# Optical Control of Cardiac Electrophysiology by the Photochromic Ligand AB2

**DOI:** 10.1101/2024.03.24.586505

**Authors:** Timm Fehrentz, Ehsan Amin, Nicole Görldt, Tobias Strasdeit, S. Erfan Moussavi-Torshizi, Philipp Leippe, Dirk Trauner, Christian Meyer, Norbert Frey, Philipp Sasse, Nikolaj Klöcker

**Author notes:** contributed equally (Timm Fehrentz, Ehsan Amin).

## Abstract

Ventricular arrhythmias (VAs) may occur in both the structurally normal and diseased heart. Particularly, patients suffering from ischemic heart disease and heart failure are at high risk of recurrent VA eventually leading to sudden cardiac death (SCD). While high-voltage shocks delivered by an implantable defibrillator may prevent SCD, these interventions themselves impair quality of life and raise both morbidity and mortality, which accentuates the need for developing novel defibrillation techniques. Here, we report the photochromic ligand azobupivacaine 2 (AB2) to enable gradual control of cardiac electrophysiology by light. By reversibly blocking voltage-gated both Na^+^ and K^+^ channels, photoswitching of AB2 modulates both the ventricular effective refractory period and conduction velocity thereby converting VA into sinus rhythm in an ex-vivo intact heart model. Thus, AB2 opens the door to the development of an optical defibrillator based on photopharmacology.

## Introduction

Ventricular arrhythmias (VAs) are life-threatening cardiac rhythm disturbances both in the structurally normal and diseased heart [66]. They comprise mono- or polymorphic ventricular tachycardia (VT) and ventricular fibrillation (VF), eventually resulting in ineffective contraction of the heart and hemodynamic compromise. The pathomechanisms underlying VA include abnormal automaticity, triggered activity, and re-entry, with the latter being predominant in patients with structural heart disease. Despite antiarrhythmic drug therapy and substantial advances in catheter ablation techniques over the past decades, VA still pose a significant risk of adverse cardiovascular events and mortality. Particularly, patients with ischemic heart disease and heart failure are at high risk of recurrent VA, worsening heart failure or even sudden cardiac death (SCD). Whereas implantable cardioverter defibrillators (ICD) may prevent SCD, there is a clear need for novel therapeutic strategies that target particularly recurrent VA, because repetitive defibrillator interventions significantly compromise quality of life and raise both morbidity and mortality themselves.

Photopharmacology allows for reversible optical control of diverse biological processes at high temporal and spatial precision [50, 71]. Photochromic ligands (PCLs) may be designed by endowing a known drug with an azobenzene moiety to afford two isomeric states of different target affinities in response to irradiation with light. PCLs have been shown to optically control ion channel activity [15, 18, 36, 44, 67] and hence physiological functions [31, 58, 72]. They hold a significant advantage over optogenetic approaches as they render gene modification of target cells dispensable when completely relying on synthetic and externally applied chromophores [4]. First attempts to develop photopharmacology towards clinical application have aimed at restoring photoreceptor function [22, 53]. Recently, we and others have applied photochromic ligands to manipulate heart rate by light [17, 42].

Here, we demonstrate that the photochromic ligand azobupivacaine 2 (AB2) allows for optical control of voltage-gated Na^+^ (Na_v_) channels. AB2 enables gradual control of cardiac electrophysiology by light, including modulation of the effective refractory period (ERP) and the conduction velocity (CV). By remote control of AB2 states, VA frequencies can be capped resulting in termination of VA and conversion into sinus rhythm (SR) in an *ex vivo* mouse heart preparation, providing proof-of-concept that AB2 or derivatives may serve as an optical defibrillator. Finally, we elucidate the biophysical principles of AB2 action in VA by mathematical modeling of cardiac tissue.

## Results

### Optical control of Na_v_1.5 channel conductance

The photochromic ligand azobupivacaine 2 (AB2) is structurally based on the voltage-gated Na^+^ (Na_v_) channel blocker bupivacaine (1), which is in clinical use as local anesthetic (Fig. 1a) [39]. The attachment of a photoresponsive azobenzene moiety created a drug existing in two states of distinct ion channel affinities (Fig. 1b). An elongated *trans*-state (2) occupied in the dark or upon illumination with blue light (480 nm) may be reversibly converted into a bent *cis*-state (3) upon UV light irradiation (385 nm). UV-Vis spectroscopy proved thermal relaxation from *cis*-AB2 to *trans*-AB2 in the dark (*k_B_T*) following a mono-exponential function with a time constant of τ = 17.6 ± 0.001 min [39]. In Fig. 1c, the principle of AB2 interaction with a voltage-gated ion channel is illustrated.

**Fig. 1.**
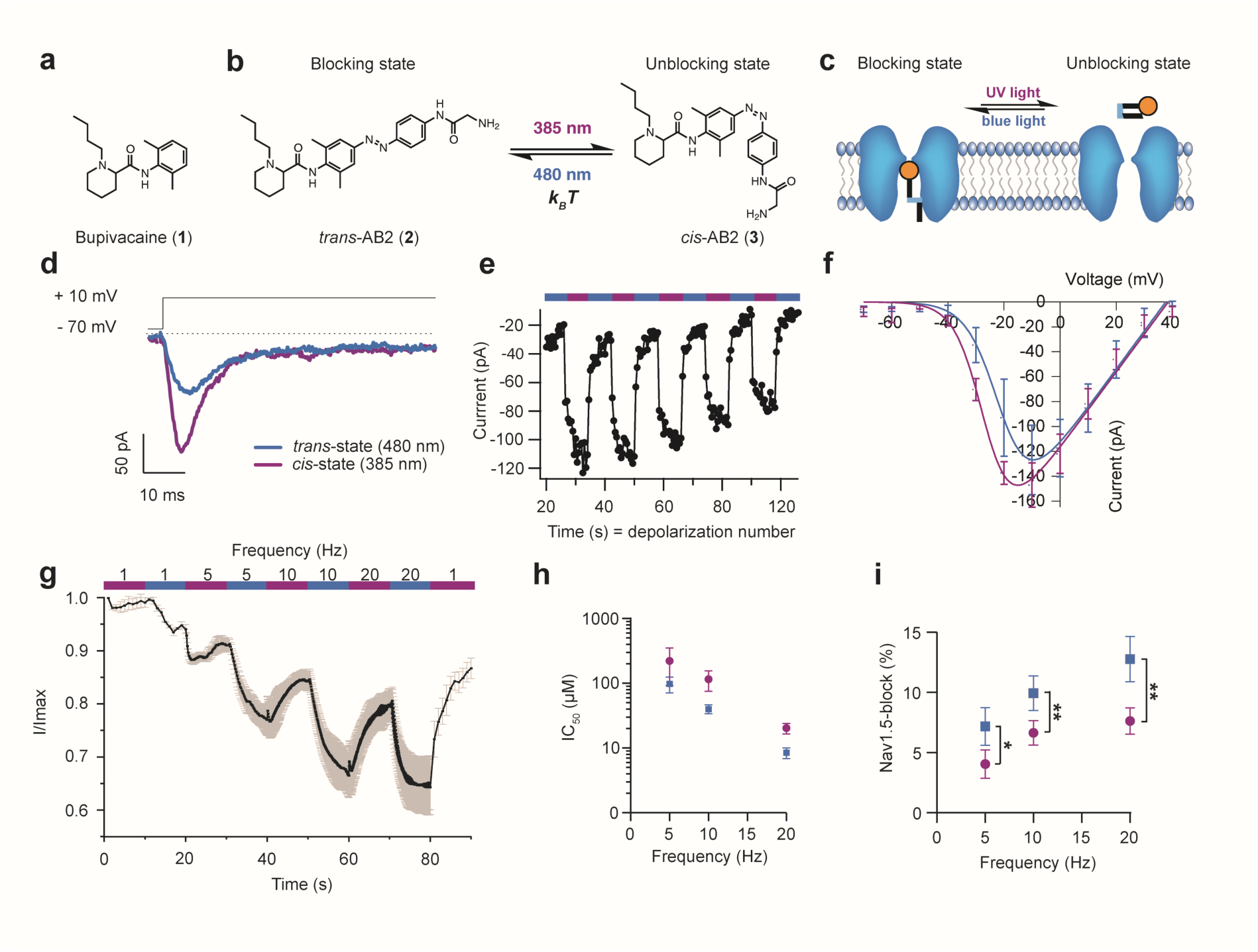
Optical control of hNa_v_1.5 channels by the photochromic ligand AB2. **a,b** Chemical structures of bupivacaine (**1**) and azobupivacaine 2 (AB2) in its *trans*- (**2**) and *cis*-state (**3**). Thermally stable *trans*-AB2 converts into *cis*-AB2 upon 385 nm light irradiation. The process may be reversed by 480 nm light irradiation or thermal relaxation in the dark (*k_B_T*). **c** Schematic illustration of AB2 acting on a voltage-gated ion channel. **d-f** Whole-cell patch-clamp recordings of the human heart Na^+^ channel hNa_v_1.5 (hH1a) stably expressed in HEK293 cells in the presence of AB2 under 385 nm or 480 nm light, respectively. Representative hNa_v_1.5 channel current traces elicited by a depolarizing voltage step in the presence of 50 µM AB2 (**d**). Na_v_1.5 channel peak currents from a representative experiment plotted over time (**e**). Current-Voltage (I-V) relationship of Na_v_1.5 channels in the presence of 50 µM AB2 in both isomeric states as indicated (**f**; n = 4 cells). **g** Normalized hNa_v_1.5 peak currents demonstrate the use-dependency of AB2 (50 µM, n = 4 oocytes). hNa_v_1.5 was activated at frequencies of 1, 5, 10 and 20 Hz under the illumination conditions indicated. **h** Use-dependency of the IC_50_ of AB2 under indicated illumination conditions (logarithmic representation of IC_50_, n = 38 oocytes). **i** Relative block of hNa_v_1.5 current by 5 µM AB2 increases with activation rate under 385 nm and 480 nm light, respectively. Color code based on respective illumination wavelengths applies to all figures. Statistical analysis: paired sample Students’ t-test (n = 6 oocytes). * p < 0.05; ** p < 0.01. All data expressed as mean ± s.e.m.

Photoswitching of AB2 was studied on hNa_v_1.5 (hH1a), the most abundant Na_v_ channel isoform in the human heart [54]. In whole cell patch-clamp experiments in HEK293 cells stably expressing hNa_v_1.5, *trans*-AB2 induced a block of inward Na^+^ currents, which was relieved when AB2 converted into its *cis*-state under UV light irradiation (Fig. 1d). The reversibility of light-dependent Na^+^ current block is shown in Fig. 1e. Quantification of AB2 photoswitching (see Methods) yielded 66.1 ± 19.2 % at a concentration of 50 µM (n = 5). The current-voltage relationship of Na_v_1.5 channels in the presence of 50 µM AB2 indicated diminished currents under 480 nm light compared to 385 nm light in the physiologically relevant voltage range (Fig. 1f). Most Na_v_ channel blockers comprising Vaughan Williams class 1 antiarrhythmic compounds exhibit a characteristic use-dependent block. This holds also true for AB2, as demonstrated in two-electrode voltage-clamp experiments in *Xenopus laevis* oocytes (Fig. 1g; n = 4). Increasing the activation frequency of hNa_v_1.5 from 1 to 20 Hz in the presence of AB2 under illumination at 385 nm and 480 nm light resulted in a use-dependent reduction of Na^+^ current. The IC_50_ of AB2 on hNa_v_1.5 channel expressed in oocytes strongly declined with increasing activation frequency from 221.9 ± 129.2 µM and 97.9 ± 27.0 µM at 5 Hz to 20.3 ± 3.8 µM and 8.4 ± 1.6 µM at 20 Hz, for 385 nm and 480 nm illumination, respectively (Fig. 1h, n = 38). These data were validated by additionally determined IC_50_ values of bupivacaine and correlation to the literature (Fig. S1). Consequently, 5 µM AB2 was sufficient to block 7.6 ± 1.1 % and 12.8 ± 4.6 % of Na^+^ current at a channel activation rate of 20 Hz, under 385 nm and 480 nm illumination, respectively. These values dropped to 4.0 ± 1.2 % and 7.2 ± 3.8 % at 5 Hz, respectively (Fig. 1i; n = 6; p = 0.0132 at 5 Hz, p = 0.0027 at 10 Hz, p = 0.0024 at 20 Hz). Together, these data show that AB2 renders hNa_v_1.5 channels light-sensitive, in addition to the previously reported block of K^+^ channels [39].

### AB2 slows signal propagation in a heart slice preparation

For a more detailed characterization of the pharmacological properties of AB2 in native tissue, we studied its effects on key determinants of cardiac excitability at first in a mouse heart slice preparation. In short-axis left ventricular slices, optical mapping of membrane voltage allowed us to describe the morphology and propagation of the cardiac action potential (AP) evoked by electrical stimulation at 5 Hz under control conditions and after superfusion with AB2 and direct radiation of the tissue slice with either UV (380 nm) or blue (480 nm) light.

As illustrated in Fig. 2a, representative activation mapping demonstrates a substantial delay in signal propagation after incubation with 5 µM *trans*-AB2 compared to control and *cis*-AB2. Representative APs from indicated regions of interest reveal extensive changes in AP morphology including a reduction in amplitude and a prolongation of AP duration in the presence of *trans*-AB2 under blue light irradiation (Fig. 2b). For the latter, we analyzed the AP duration at 80 % repolarization (APD_80_) as both the human APD_75_ - APD_85_ [5, 7, 20, 38, 55] and the mouse APD_80_ [34] are well-known surrogates of the tissue specific ventricular effective refractory period (ERP). Quantification of data yielded a reduction in conduction velocity (CV) by 7 % and 33 % (p = 0.6, p = 0.004; n = 5; Table 1, Fig. 2c), a prolongation of the APD_80_ by 10 % and 46 % (p = 0.5, p = 0.0003; n = 5; Table 1, Fig. 2d), and a decrease in AP amplitude by 4 % and 27 % (p = 0.8, p = 0.005; n = 5; Table 1, Fig. 2e) in *cis*-AB2 and *trans*-AB2, respectively, compared to control conditions. As AP amplitude, repolarization, and propagation rely considerably on the voltage-dependent interplay of Na^+^ and K^+^ channels, these observations are in good agreement with *trans*-AB2 blocking both Na_v_ and K_v_ channels, hence raising the level of our previous recombinant experimental data to native cardiac tissue.

**Fig. 2.**
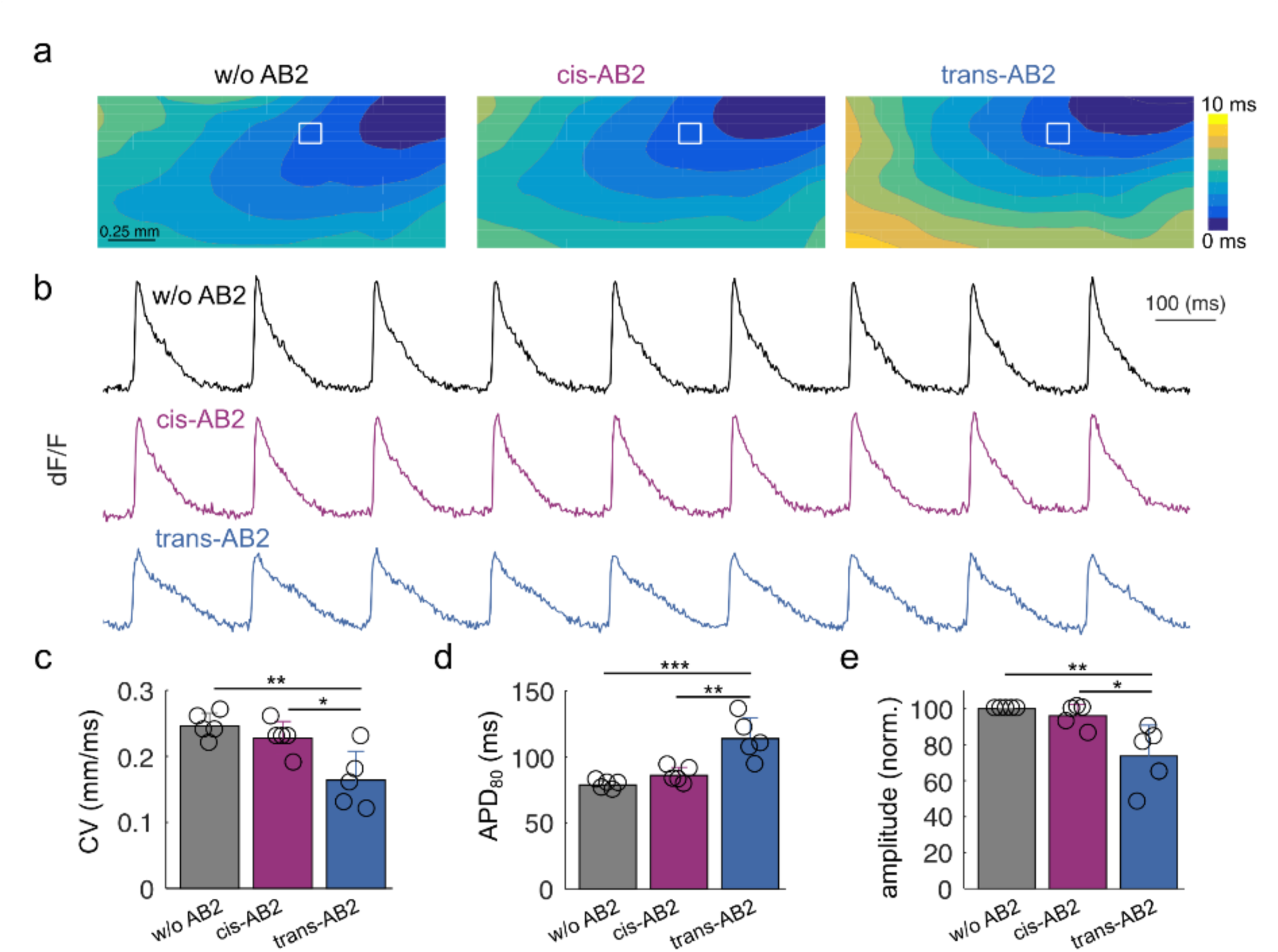
AB2 reduces cardiac conduction velocity (CV) and reshapes the cardiac action potential (AP) in mouse left ventricular slices in a state-dependent manner. **a** Representative activation time maps of cardiac tissue under control conditions without AB2 (w/o AB2) and after application of 5 µM under UV (*cis*-AB2) and blue (*trans*-AB2) light in a mouse heart slice preparation. Isochrones distances are 1 ms. **b** Representative APs from indicated regions of interests (white squares in a) are shown under control, *cis*-AB2, and *trans*-AB2 conditions. AP amplitudes are presented as relative change in fluorescence intensity (ΔF/F in %). **c - e** compared to *cis*-AB2 and w/o AB2, *trans*-AB2 significantly reduces CV (c), prolongs APD_80_ (d), and simultaneously reduces AP amplitude (e). Statistical analysis: one-way ANOVA and posthoc analysis with Tukey-Kramer test (n = 5); * p ≤ 0.05; ** p ≤ 0.01; *** p ≤ 0.001. All data are expressed as mean ± s.e.m.

**Table 1.** Summary of changes in conduction velocity (CV), action potential duration (APD_80_), and amplitude by AB2 in a state-dependent manner.

|  | CV (mm/ms) | APD<sub>80</sub> (ms) | rel AP amplitude (%) |
| --- | --- | --- | --- |
| w/o AB2 | 0.246 ± 0.019 | 78 ± 3 | 100 |
| cis-AB2 | 0.228 ± 0.025 | 86 ± 6 | 96 ± 6 |
| trans-AB2 | 0.164 ± 0.044 | 114 ± 16 | 73 ± 17 |

### Isomer-dependent effects of AB2 on cardiac conduction and field potentials in whole heart tissue

Na_v_ channel blockers slow cardiac CV by reducing the rate of initial depolarization of the cardiac AP [26, 33]. In the human electrocardiogram, broadening of the QRS complex is pathognomonic. We therefore assessed how AB2 affects spontaneous epicardial field potentials (FP) recorded as bipolar cardiac electrograms from a Langendorff *ex vivo* mouse heart preparation. AB2 was applied within the perfusate and converted into its *cis*- or *trans*- state by pre-irradiation with 385 nm or 480 nm light prior to entering the coronary arteries. Fig. 3a shows a representative bipolar FP under control conditions. Upon perfusion of the heart with *cis*-AB2 (10 µM), slight FP broadening occurred (Fig. 3b), which was, however, much more pronounced when AB2 was perfused in its *trans*-state (Fig. 3c). The temporal course of FP broadening under control, *cis*-AB2, and trans-AB2 is shown in Fig. 3d for a representative experiment. The observed increases in the slope of spontaneous FP width upon *cis*- and *trans*-AB2 application, respectively, are quantified in Fig. 3e. The slope increased from control to *cis*-AB2 by a factor of 6.34 ± 2.8 (n = 5), which was insignificant to the control situation (p = 0.12). However, upon application of *trans*-AB2, the slope increased significantly as much as 23.1 ± 7.4 fold (n = 5; p = 0.037 compared to *cis*-AB2). In addition, we paced the ventricle and recorded evoked FP (6.6 Hz, 400 bpm). Representative FP under control and indicated AB2 conditions are shown in Fig. 3f-h. Both the latency and width of the evoked FPs increased by application of AB2 over time in a state-dependent manner (Fig. 3i, Fig. S2). The slope of the increase in latency over time steepened significantly by 6.9 ± 2.0 fold after application of *cis*-AB2, whereas *trans*-AB2 increased it by a factor of even 49.5 ± 12.8. (Fig. 3j, n = 4). The slope of the increase in stimulated FP width rose from control to *cis*-AB2 by a factor of 7.2 ± 1.0, whereas *trans*-AB2 increased it by even 72.3 ± 23.9 fold (Fig. S3, n = 3).

**Fig. 3.**
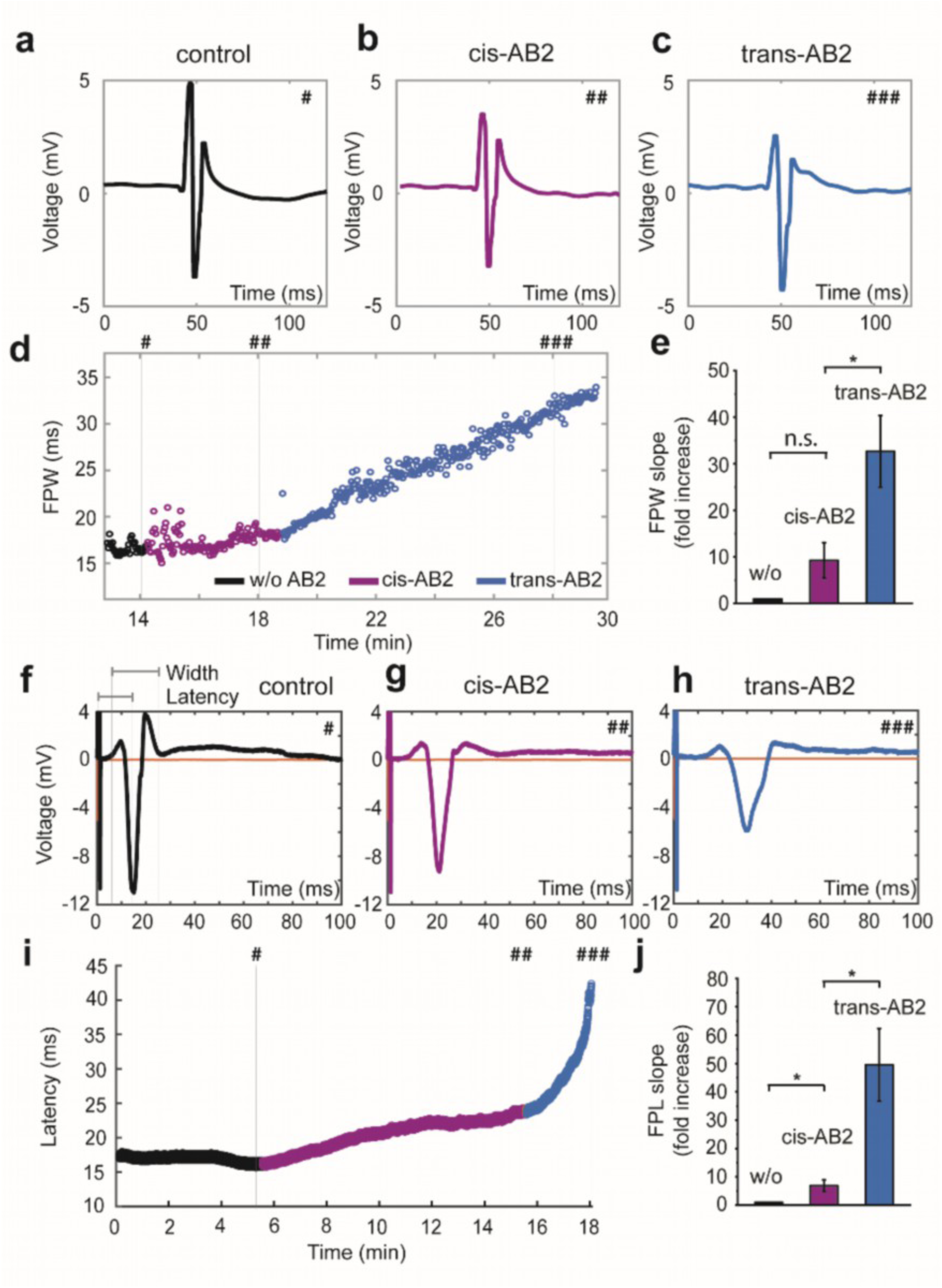
AB2 slows spontaneous and evoked FPs in a Langendorff-perfused mouse heart preparation. **a-e** Effects of AB2 on the duration of spontaneous FPs. Representative FPs under control conditions without (w/o) AB2 (**a**) and in the presence of 10 µM AB2 in its *cis*- and *trans*-state, respectively (**b**, **c**). Representative temporal course of FP broadening (FP width, FPW) under control, *cis*-AB2, and *trans*-AB2 conditions as indicated (**d**) and its quantification (**e**; paired t-test, w/o AB2 to *cis*-AB2 p = 0.12, *cis*-AB2 to *trans*-AB2 p = 0.037, n = 5 hearts). Rhombus correlate FP from a-c with their time points in d. Color code of d applies for all figures. **f-j,** Effects of AB2 on the latency of ventricular FPs evoked by electrical pacing at 6.6 Hz (400 bpm). Representative ventricular FPs under control conditions without (w/o) AB2 (**f**) and in the presence of 10 µM AB2 in its *cis*- and *trans*-state, respectively (**g**, **h**). Representative temporal development of the FP latency under control, *cis*-AB2, and *trans*-AB2 conditions as indicated (**i**) and its quantification (**j**; paired t-test, w/o AB2 to *cis*-AB2 p = 0.0094, *cis*-AB2 to t*rans*-AB2 p = 0.0034, n = 4 hearts). Hash marks correlate FPs from **f-h** with their time points in **i**. n.s. = not significant, * p < 0.05. All data expressed as mean ± s.e.m.

These data obtained from a whole heart preparation demonstrate that AB2 slows both spontaneous and evoked bipolar FPs, distinctly depending on its isomeric state: in its *trans*- state, AB2 was much more effective in slowing ventricular conduction than in its *cis*-state.

### Isomer-dependent effects of AB2 on cardiac excitability in whole hearts

Exploiting AB2 for gradual control of cardiac excitability and its development into an optical defibrillator would require *cis*-AB2 having either no or negligible effects on the effective refractory period (ERP) and CV in order not to disrupt physiological signal propagation. To find robust experimental conditions, we screened AB2 concentrations for its effects on ventricular ERP as assessed by programmed electrophysiological stimulation in the Langendorff-perfused *ex vivo* mouse heart applying a conventional S1-S2 protocol (Fig. 4a). In parallel, we monitored changes in CV under a basic cycle length (BCL) of 150 ms. It should be noted, that control ERP values varied considerably from one individual to another, which is in good agreement with the literature reporting a broad range of ERP values from 33 ms up to 80 ms for mouse ventricular tissue [32].

**Fig. 4.**
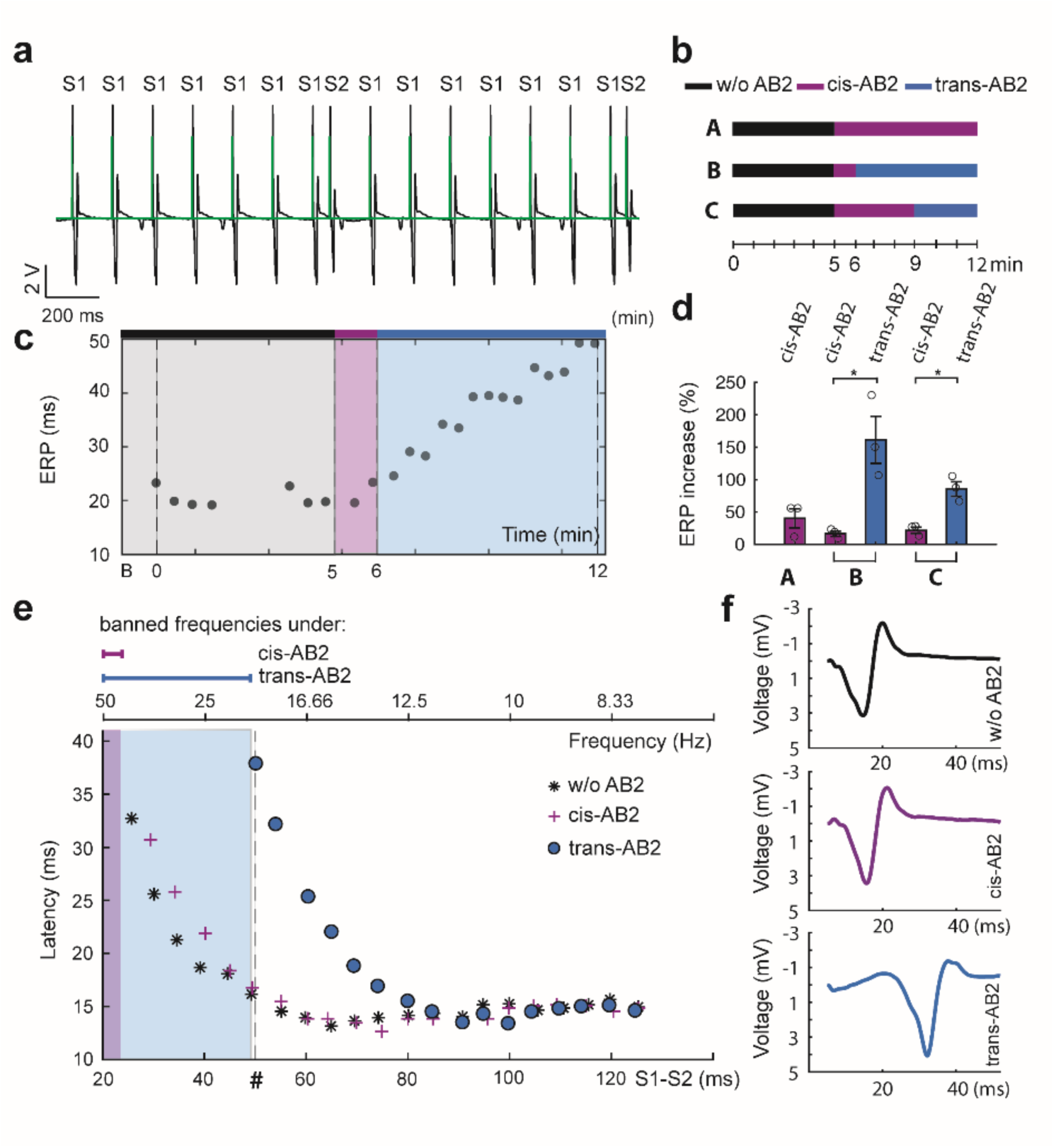
AB2 prolongs ERP in a Langendorff-perfused mouse heart preparation. **a** Electrophysiological stimulation followed a canonical S1-S2 stimulation protocol, with the stimuli being indicated in green and the corresponding field potentials (FPs) in black. Electrical stimulation was applied to the right ventricle and the resulting FP was detected at the left ventricle. The protocol comprises consecutive trains of six S1-S1 intervals (basic cycle length) of 150 ms length (6.6 Hz), followed by a S1-S2 interval shortening every iteration by 1, 3, or 5 ms starting from 150 ms. At the end of the protocol, a S2-S1 interval of 150 ms followed. The resulting FPs within the S2-S1 interval were analyzed for latency and refractory period. The first electrical stimulus that fails to produce a FP during the S2-S1 interval represents the effective refractory period (ERP), which correlates with a distinct excitation frequency. **b** Protocols A - C of AB2 application: protocol A includes 5 min of control (w/o AB2) followed by 7 min of *cis*-AB2 application. Protocol B comprises 5 min control (w/o AB2), 1 min of *cis*-AB2 and 6 min of *trans*-AB2. In protocol C, 5 min of control (w/o AB2) are followed by 4 min of *cis*-AB2 and 3 min of *trans*-AB2. Color code applies for all figures. **c** Representative time course of changes in ERP under protocol B conditions at 5 µM AB2. **d** Quantification of normalized ERP increase under *cis*- and *trans*-AB2 compared to control according to protocols A to C (n = 3 hearts, respectively). **e** Representative FP latencies plotted over S1-S2 intervals following protocol B. Excitation frequencies banned by AB2-mediated ERP prolongation are indicated in color-code. **f** Single representative FPs from Fig. 4e at the position indicated by the rhombus (S1-S2 = 50 ms) showing an increase in latency under *trans*-AB2. Statistical analysis: paired sample Students’ t-test (n = 3). * p < 0.05. All data expressed as mean ± s.e.m.

We identified a concentration of 5 µM AB2 to satisfy at its best the above conditions under low [K^+^]_e_ (2 mM). After 10 min of perfusion, 5 µM *cis*-AB2 raised the control ERP (w/o AB2) from 30.7 ± 6.4 ms to 44.8 ± 4.3 ms (Fig. S3, n = 3), whereas CV decreased by 5.9 ± 3.3 % (Fig. S5). A steady-state was reached already within the first 3 min of application. Subsequent perfusion with *trans*-AB2 for only 4 min doubled approximately ERP up to 89.0 ± 9.3 ms (Fig. S4, n = 3), whereas CV was reduced by 13.0 ± 7.5 % (Fig. S5), which is insignificant from the above reduction in CV induced by *cis*-state AB2 (Fig. S5, paired t-test, p = 0.2984, n = 3 hearts).

Based on this preliminary work, we designed three different experimental protocols, each lasting a total of 12 min (A - C; Fig. 4b): After an initial period of S1-S2 stimulation for 5 min to determine ERP under control conditions, *cis*-AB2 and *trans*-AB2 were applied for indicated times, allowing a more detailed analysis of the AB2 efficiency profile.

Following protocol A, the average ERP was 30.7 ± 6.4 ms under control conditions. Subsequent application of *cis*-AB2 for 7 min increased the ERP by 40.2 ± 14.8 % yielding 41.3 ± 5.2 ms (Fig. 4d, Fig. S4, n = 3). The ERP observed following protocol A would thus theoretically allow maximum ventricular excitation frequencies of 33 Hz under control and of 24 Hz under *cis*-AB2 conditions. CV was reduced by only 7.5 ± 2.6 % (Fig. S5, see Methods).

Following protocol B, the control ERP increased from 18.6 ± 2.0 ms to 24.8 ± 1.6 ms after 1 min of *cis*-AB2 perfusion, equaling an increase by 17.0 ± 4.6 % (Fig. 4d, n = 3). Under the subsequent *trans*-AB2 conditions for 6 min, ERP rose to 55.8 ± 9.4 ms resembling an increase by 161.4 ± 36.2 % at the end of the 12 min protocol (Fig. 3d, paired t-test, p = 0.045, n = 3 hearts). The ERP observed following protocol B would thus theoretically allow maximum ventricular excitation frequencies of 54 Hz under control, of 40 Hz under *cis*-AB2, but only 18 Hz under *trans*-AB2 conditions. CV increased by 0.7 ± 0.7 % after *cis*-AB2 application, whereas *trans*-AB2 reduced CV by 8.0 ± 3.3 %. However, both changes in CV were statistically insignificant from each other (Fig. S5, paired t-test, p = 0.13, n = 3 hearts). Protocol C differs from protocol B in a prolonged *cis*-AB2 application. It was designed to closer mimic a future *in vivo* situation, in which AB2 would be present all the time – either in its *cis*- or its *trans*-state. Average ERP increased from 26.9 ± 6.0 ms under control to 32.2 ± 5.8 ms under *cis*-AB2, and to 48.6 ± 7.9 ms under *trans*-AB2 conditions (Fig. S5, n = 3). The shift to *trans-*AB2, but not to *cis*-AB2, represents a significant increase over control (Fig. 4d, paired-sample t-test, p = 0.013). These data translate into relative increases by 21.8 ± 4.8 % under *cis*-AB2 and by 85 ± 11.2 % under *trans*-AB2 conditions. For protocol C, the observed ERP would theoretically allow for maximum ventricular excitation frequencies of approximately 37 Hz under control, 31 Hz under *cis*-AB2, and only 21 Hz under *trans*-AB2 conditions. CV was reduced by 6.3 ± 1.6 % and 11.3 ± 1.7 % after *cis*- and *trans*-AB2 application, respectively. Both changes were statistically insignificant from each other (Fig. S5, paired t-test, p = 0.25, n = 3 hearts). A comprehensive quantification of the state-dependent ERP prolongation and CV changes by AB2 is given in Fig. 4d and Fig. S5.

Finally, the latencies of FPs evoked by S2 from the respective last S1-S2 stimulation following protocol B were plotted against the corresponding S1-S2 pacing intervals. As shown in Fig. 4e, the characteristic increase in latency upon shortening the S1-S2 intervals was preserved after application of AB2, independent of its isomeric state. However, *trans*- AB2 increased FP latencies at given S1-S2 intervals to much greater extent than did *cis*- AB2, which was hardly distinct from untreated control. Banned ventricular excitation frequencies resulting from the above-described prolongation of ERP by *cis*- and *trans*-AB2, respectively, are indicated in color-shaded background. Representative S2-S1 FPs under indicated experimental conditions are depicted in Fig. 3f.

Together, the shift between the two isomeric states of AB2 adjusts the maximum ventricular excitation frequencies, creating a range for precise control of cardiac excitability. Turning *cis-* into *trans-*AB2 bans excitation frequencies > 18 Hz following protocol B and > 21 Hz following protocol C, respectively, and may thereby terminate VA.

### Isomer-dependent defibrillation of VA by AB2

Finally, we explored photoswitching of AB2 as a potential means to terminate VA. Stable runs of VA were induced in the Langendorff-perfused mouse heart by either low [K^+^]_e_ of 2 mM alone or combined with electrical burst or S1-S2 stimulation (see Methods) as shown before [10, 11, 16, 21, 65]. VAs were considered stable when they lasted > 5 min without spontaneous termination. In addition to control experiments in the absence of AB2 (w/o AB2), the above-described protocols of 12 min were used again for applying the respective isomers of AB2 (Fig. 4b). Fig. 5 illustrates representative epicardial electrogram recordings of a stable VA, which was successfully terminated by photoswitching AB2 following protocol B (Fig. 5a - c). Frequency spectra were extracted from the respective electrogram recordings by Fourier analysis (Fig. 5d - f) indicating a representative shift from frequency bands of > 20 Hz under control conditions to < 5 Hz under *trans*-AB2 conditions. For statistical analysis of VA termination, the last 6 minutes of protocols A, B, and control situation (w/o AB2) were compared. Termination was categorized to be either successful or not. VAs were considered successfully terminated by the appearance of single FP reflecting QRS complexes that consistently transitioned into sinus rhythm (SR). For more detailed information, individual FP recordings at the end of protocols A and B (00:11:40 - 00:11:50, hour: min: sec) are shown in Fig. S6.

**Fig. 5.**
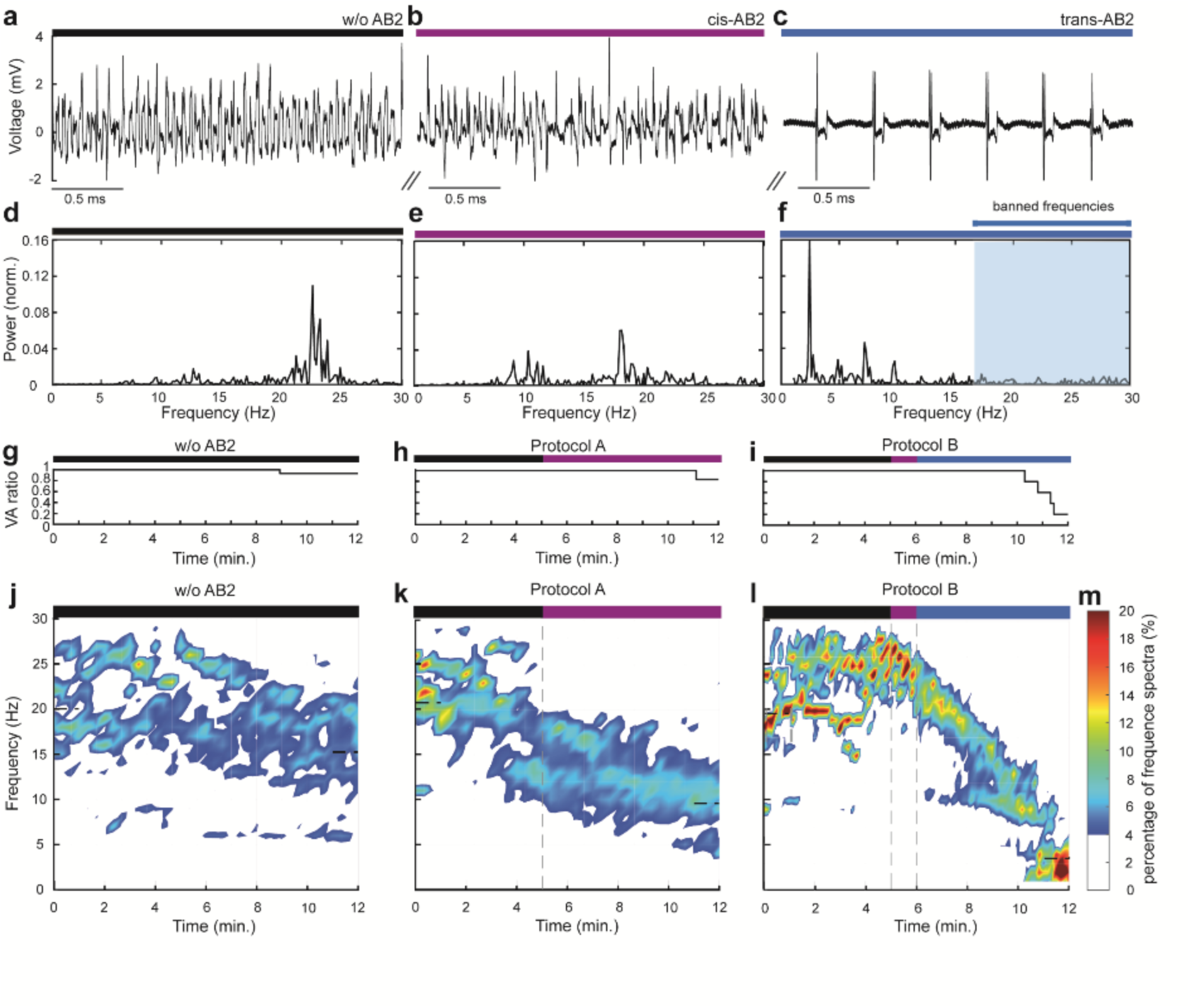
Termination of ventricular tachycardia by state dependent AB2 application. **a-f** Representative experiment of terminating a ventricular arrhythmia (VA, including VT and VF) by AB2 in the Langendorff-perfused mouse heart following protocol sequence B (Fig. 4b). **a** Cardiac electrogram showing a stably running VA in the absence of AB2 at the end of the 5 min period. **d** Fourier analysis of VA frequency spectrum found in a. **b** Cardiac electrogram 1 min after application of 5 µM *cis*-AB2. **e** Fourier analysis of VA frequencies found in b. Note that the dominant frequencies are shifted towards lower frequencies. **c** Sinus rhythm (SR) 3 min after optical defibrillation of the VA by *trans*-AB2. **f** Fourier analysis of SR frequency spectra found in c**. g-I** VA maintenance under control condition (n = 15 hearts) and protocols A (n = 6 hearts) – B (n = 5 hearts) over 12 min in alignment to **j-l**, respectively. **j** Heat map of normalized control VA frequencies according to control measurements (n = 15 hearts). Frequencies below 1 Hz are not shown. **k** Heat map of normalized VA frequencies under AB2 application according to the color-coded protocol sequence A (n = 6 hearts). **l** Heat map of normalized VA frequencies under AB2 application according to the color-coded protocol sequence B (n = 5 hearts). **m** Color map of the frequency percentage at a given time interval applies to (**j**-**l**).

Altogether, one out of 15 hearts (6.7 %) of VA terminated under control situation (w/o AB2; n = 15 hearts). *Cis*-AB2 application following protocol A terminated 1 out of 6 hearts (16.7 %) of VA (n = 6 hearts). However, *trans*-AB2 terminated 4 out of 5 hearts (80 %) of running VA following protocol B (n = 5 hearts). Under protocol C, *trans*-AB2 application terminated 2 out of 3 hearts (66.6 %) of still running VA within the last 3 minutes (n = 3 hearts). VA maintenance of protocols A, B, and control (w/o AB2) over 12 min are plotted in Fig. 5 g-I. The last 6 min were analyzed by the cox regression [60, 61] and represented as survival curves Kaplan Meier (Fig. S7). Comparison of control with *cis*-AB2 (protocol A) revealed no significant difference (score (logrank) test = 0.41, p = 0.5). However, significant differences were found between *cis*-AB2 (protocol A) and *trans*-AB2 (protocol B) (Score (logrank) test = 4.0, p = 0.04), and even more so between control and *trans*-AB2 (protocol B) (score (logrank) test = 10.8, p = 0.001). The average duration for terminating respective VA in our experimental setup was 00:04:48.75 ± 44.5 (hour: min: sec) after *trans-AB2* application in protocol B and 00:02:27.5 ± 44.5 after *trans*-AB2 in protocol C.

Data were also analyzed for the temporal changes in VA frequency spectra during the complete protocol length of 12 min. In Fig. 5 j-m, heat maps of normalized VA frequency spectra for indicated protocol conditions are depicted. Weighted frequencies at the first and the last minute of respective protocols were calculated (see Methods): In the absence of AB2 (w/o AB2), the weighted frequency decreased from 19.9 ± 4.5 Hz to 15.2 ± 5.3 Hz (Fig. 5j; n = 15), with all but one VA still ongoing at the end of the 12 minutes lasting protocol. In part, we attribute the reduction in VA frequency to a decrease in amplitude consequently challenging the automated detection algorithm. However, weighted VA frequencies within the first minute were in good agreement with values ranging from 15 - 30 Hz as reported in the literature [16].

Under protocol A, *cis*-AB2 decreased the weighted frequency more strongly from 20.91 ± 4.4 Hz to 9.5 ± 3.2 Hz, albeit the VA still persisted and spontaneous SR was detected again in only a single heart (Fig. 5k, Fig. S6, n = 6). When following protocol B, however, *trans*-AB2 reliably shifted weighted frequencies from 19.0 ± 5.3 Hz all the way down to 3.6 ± 2.1 Hz and eventually terminated VA (Fig. 5l, n = 5). Four out of five hearts showed termination of VA with a consistent transition into SR (Fig. S6, n = 5), whereas only a single heart remained in a VA. It is important to note that the frequencies of SR in Langendorff-perfused mouse hearts ranged between 5 - 7 Hz, which is in good agreement to the reported literature [16]. AB2 may hence prove useful not only to limit VA frequencies but also to effectively terminate stably running VA.

## Discussion

Malignant cardiac arrhythmias like VT and VF represent life-threatening conditions that may require defibrillation. Yet, defibrillator interventions by high-voltage shocks have severe side effects including pain, anxiety, subsequent depression as well as myocardial tissue damage increasing mortality when performed repetitively [2, 46, 48]. Since common pharmacological approaches and catheter ablation techniques have proven only limited success in preventing VA, there is still an urgent medical need for developing novel strategies to prevent and treat VA [19, 41, 69]. In the present study, we introduce AB2 as an innovative tool for gradually controlling cardiac excitability by light that may eventually open a new avenue towards clinical application of optical defibrillation.

The envisioned optical approach implies a PCL adopting two isomeric states of distinct receptor affinities, which may be switched by light of different wavelengths. AB2 meets such requirements and allows for reliable optical control of Na_v_1.5 channel function [57]. Whereas the bent *cis*-state of AB2 reduces Na^+^ currents rather moderately, *trans*-AB2 is significantly more effective in blocking them. The block is reversible and occurs throughout the physiologically relevant range of voltage-dependent channel activation. As known for other Na^+^ channel blockers derived from the family of local anesthetics, AB2 blocks Na_v_1.5 in a use-dependent manner with channel gating being a requisite for the drug to bind [29]. However, the difference in state dependent block by *cis-* vs. *trans*-AB2 is preserved for frequencies up to 20 Hz. Thus, Na_v_1.5 inhibition by AB2 strongly increases with the frequency of channel activation with decreasing IC_50_ values. *Trans*-AB2 does not exhibit the same efficacy as the parent compound bupivacaine, indicating that attaching an azobenzene moiety reduces its apparent affinity to some extent. This has also been observed in similar studies for FHU-779, an L-type Ca^2+^ channel PCL derived from diltiazem [17]. Nevertheless, AB2 shares important pharmacological properties with bupivacaine, from which it is structurally derived. Besides blocking Na_v_1.5 channels, bupivacaine also blocks most cardiac K_V_ channels including members of the K_v_1, K_v_2, K_v_4, K_v_7, and K_v_11 subfamilies at IC_50_ values in the lower µM range [24, 25, 40, 59], reducing the repolarizing currents I_to_, I_Kur_, I_Kr_, and I_Ks_ of the cardiac AP. This is in good agreement with the data we obtained for AB2 lowering AP amplitude and prolonging APD_80_ in optical mapping experiments on acute mouse heart slices. In addition, the delay in signal propagation by AB2 may well be the result from Na_v_1.5 block, as CV relies, at least in part, on the initial depolarization rate of the AP [33]. Thus, the effects of AB2 on cardiac electrophysiology that we demonstrate for acute cardiac slices and ex vivo whole heart preparations are most likely the result of a combined block of both Na_v_ and K_v_ channels.

Na_v_ channel blockers like class 1 antiarrhythmic drugs and bupivacaine slow ventricular CV, which may be revealed by QRS broadening in the human electrocardiogram [27]. However, the correlation between QRS width, reflecting the time of electrical activation of the ventricles, and CV might be more complex in mice [9]. Here, we first recorded spontaneously generated FPs to characterize physiological signal propagation. In a bipolar recording mode, FP width correlates inversely with CV [3]. Determining the latency of evoked FPs complemented our estimate of CV. AB2 broadened both spontaneous and evoked FPs in a state-dependent manner. Also, the latency of evoked FPs increased after application of *trans*-AB2 as compared to *cis*-AB2. Thus, the observed effects of AB2 on murine bipolar cardiac electrograms extend our findings in heart tissue slices and strongly suggest that besides slowing CV, AB2 also decelerates ventricular signal propagation.

The ERP estimates the excitability of cardiac tissue and defines a maximum activation frequency. As such, the ERP in combination with the CV may be considered as an indicator of VT and VF stability. In the present study, we determined the ERP in a whole heart preparation using S1-S2 protocols that were previously applied in both mouse and human ventricular myocardium [51, 56]. In good agreement with the recorded FPs described above, the state-dependency of AB2 effects held also true for the observed increase in ERP at all concentrations applied. In an orienting screen, we identified a concentration of 5 µM at which *cis*-AB2 showed only moderate effects, whereas *trans*-AB2 prolonged ERP sufficiently to prevent or even terminate VA. It is essential to realize that the increase in ERP by AB2 being shifted from its *cis*- into *trans*-state severely restricts ventricular activation frequencies under reduced CV. The consequent ban of high frequencies may destabilize and terminate VT and VF. ERP correlates well with the APD at the cellular level and is dominantly influenced by repolarizing K^+^ conductances. However, in wild type mouse heart, reported ERP values vary greatly between 33 and 80 ms [32]. In a heterozygous Scn5a^+/-^ mouse model, ventricular ERP was determined to be 56 ± 7 ms, approximately twice as long as in their wild type littermates [51], which compares well to the AB2 effects observed in the present study. Altogether, it remains unclear to what extent Na_v_ or K^+^ channel blocking properties of AB2 explain its prolongation of ventricular ERP.

Several theoretical concepts have been developed to deeper understand atrial and ventricular arrhythmia including the leading circle theory [1, 12] and the more sophisticated and generally applicable spiral wave theory [12, 49]. According to the leading circle theory, the cardiac wavelength (λ = ERP x CV) determines a minimum path length for re-entry and thus predicts whether the arrhythmia will be stable or not under given conditions. An extended cardiac wavelength diminishes the likelihood of reentry and ultimately exhibits antiarrhythmic effects. This theory resulted from studies of atrial arrhythmias and offers a simplified approach to deduce the mechanism of action of antiarrhythmic drugs, particularly effective in treating atrial fibrillation. For instance, typical class III antiarrhythmics like sotalol and amiodarone, which block predominantly K^+^ and to lesser extent Na_v_ channels, destabilize reentry arrhythmias [28] by increasing APD and consecutively ERP [5, 7, 20, 34, 38, 55]. However, class Ic antiarrhythmics including flecainide, which target predominantly Na_v_ channels, do not necessarily increase the cardiac wavelength but may still terminate re- entry arrhythmias [70]. Such findings have challenged the original leading circle theory and gave rise to considerations summarized by the spiral wave theory [12], which describes both atrial and ventricular fibrillation [49]. By optical mapping of membrane voltage in an acute heart slice preparation, we found AB2 to both significantly decrease CV and prolong APD, whereas electrophysiological stimulation of whole heart tissue revealed a predominant increase in ERP by AB2, which was accompanied by only a rather mild reduction in CV. Such different findings may reflect differences between the two models, including tissue structure and recording methodology. Taking also our recombinant electrophysiological data into account, we assume that AB2 acts on both Na_v_1.5 and K^+^ channels at relevant concentrations to terminate VA, irrespective of whether the leading cycle or the spiral wave theory may be more suitable to describe our data. Still, to get a more detailed understanding of VA termination by AB2, mathematical modeling was performed. Fitting the most characteristic changes in AP morphology, which we had observed by optical mapping after application of *trans*-AB2, indicated a central role for I_Nav_ and I_Kur_ inhibition (Fig. S8). In a 2D cardiac tissue model, only simultaneous inhibition of I_Nav_ and I_Kur_ reliably terminated the otherwise sustained VA after non-linear meandering and eventual collision of the spiral cores (Fig. S9), strongly supporting our interpretation of the experimental data. On a perspective, optical mapping of VA and their photopharmacological termination in the whole heart preparation will be helpful for improving our mechanistic understanding of AB2 action.

Whereas strategies like anti-tachycardia pacing [19, 41, 69] have been developed in addition to electrical defibrillation by high-voltage shocks during recent years, first attempts to develop optical defibrillations have also been reported. They were all based on optogenetic tools: cardiomyocytes were transduced with the membrane depolarizing light-gated ion channel ChR2 or the membrane hyperpolarizing proton pump ArchT [10, 14, 21, 47, 63] enabling termination of VA upon direct epicardial illumination. By contrast, our photopharmacological approach holds the advantage of avoiding any potential risk associated with genetic manipulation of human tissue. To improve the functionality of AB2, sign-inversion in action might be useful: by exchanging the azobenzene by a diazozine moiety, AB2 may be turned into a *cis-state blocker* [62]. However, the presented photopharmacological approach seems currently too slow to be feasible in a clinical setting. For our experiments, we interpret the slow time course of VA termination as caused by the slow diffusion of AB2, photoswitched a priori, into the tissue. To overcome this limitation, the azobenzene absorption spectrum in AB2 should be shifted to longer wavelengths [35], enabling photoswitching AB2 more deeply within the tissue and hence directly at its receptor binding site. Such an approach would likely significantly accelerate drug action. Besides application in optical defibrillation, the photochromic ligand AB2 generally offers the great chance for controlling cardiac excitability at a temporal and spatial precision by light, which may well complement clinical diagnostics and therapy in the heart electrophysiology lab. After our first proof of concept application of the photochromic diltiazem derivative FHU-779 for optical modulation of heart rate, the present study significantly expands the application spectrum of cardiac photopharmacology.

## Acknowledgements

The authors thank Hugues Abriel (University of Bern) and Robert Kass for the pcDNA 3.1 vector encoding α-subunit of hNa_v_1.5 (Addgene plasmid #145374) and Hugues Abriel for providing the HEK293 cell line stably expressing hNa_v_1.5. Furthermore, we would like to thank Stefan Schaetz for valuable technical support. N.K. received support from the Deutsche Forschungsgemeinschaft (SFB 1116, TP A01). P.S. was supported by the Deutsche Forschungsgemeinschaft (GRK1873, SA 1785/7-1, SA 1785/9-1).

## Author contributions

The project was conceived by T.F., N.K. and P.S. Patch clamp characterization of AB2 was carried out by N.G. and T.F. Two-electrode voltage-clamp recordings in *Xenopus laevis* oocytes were performed by T.S. Optical mapping experiments were performed by E.A. and S.E.M. Langendorff-perfused heart experiments were performed by T.F. together with N.G.. Data analysis from Langendorff-perfused heart experiments was carried out by E.A. and T.F.. P.L. advised on photoswitch choice and synthesized AB2. Mathematical modeling was performed by E.A.. P.S., N.F., D.T. and C.M. helped to discuss the results. T.F. and N.K. wrote the manuscript with contributions from all authors.

## Competing interests

The authors declare no competing interests.

## Materials and Methods

### General

Animals were housed in a central animal facility of the University of Düsseldorf. All animal experiments were carried out according to the Directive 2010/63/EU of the European Parliament on the protection of animals used for scientific purposes, which is congruent to the German animal welfare act and the laboratory animal welfare ordinance. Furthermore, all procedures were approved by the responsible animal ethics committee (State Agency for Nature, Environment, and Consumer Protection, LANUV).

### Cell Culture

A stable HEK293 cell line expressing Na_v_1.5 (tested for mycoplasma, not authenticated) was cultured in Dulbeccós modified Eagle medium (DMEM, GiBCO) containing 10 % fetal bovine serum (FBS, Biochrome) in an incubator at a 37 °C and 5 % CO_2_. Cells were splitted at ∼ 80 % confluency by applying trypsin. Cells were seeded on treated glas coverslips at a density of ∼ 13000 cells per cm^2^. Patch clamp experiments on cells were performed 16-25 h after splitting.

### *In vitro* transcription and expression of hNa_v_1.5 in Xenopus oocytes

Prior to *in vitro* transcription, a hNa_v_1.5 plasmid (Addgene plasmid #145374; http://n2t.net/addgene:145374; RRID:Addgene_145374) was linearized with XbaI. For synthesis of complementary RNA (cRNA), mMESSAGE mMACHINE ® T7 Transcription Kit (ThermoFisher Scientific, Waltham, MA, USA) was used. The quality of synthesized cRNA was checked by 1% agarose gel electrophoresis. Stage IV and V *Xenopus* oocytes were purchased from EcoCyte Bioscience (Dortmund, Germany) and injected with 25 ng of Na_v_1.5 cRNA per oocyte. For injection, a nanoliter injector was used (SYS-Micro4, WPI, Sarasota, FL, USA). Oocytes were cultured at 17°C in Barth’s medium, containing (in mM): 88 NaCl, 2.4 NaHCO_3_, 1 KCl, 0.8 MgSO_4_, 0.4 CaCl_2_, 0.33 Ca(NO_3_)_2_ and 20 HEPES (titrated to pH 7.5 with NaOH). The medium was supplemented with 100 units/mL penicillin, 0.1 mg/mL streptomycin and 0.25 µg/mL amphotericin B (antibiotic antimycotic solution, Sigma Aldrich, St. Louis, MO, USA). Voltage clamp recordings were performed after 1-2 days of expression.

### Electrophysiology

Patch clamp recordings of whole cell Na_v_1.5 currents in HEK293 cells were performed using a HEKA Patch Clamp EPC10 USB amplifier and patch master software (V2X90, HEKA) as described before [17]. The resistance of patch pipettes (Science Products) ranged between 3.5 - 5 MΩ. Illumination of samples was carried out by using a custom-built light source [17], including high power UV-A (385 nm) and blue (470 nm) LEDs (Thorlabs), resulting in approximately 32.8 mW/mm^2^ and 32.1 mW/mm^2^ light intensity in the field of view, respectively (water immersion objective, Olympus, LUMPLFL N 40x, field number 26.5, NA 0.8). LED light was coupled into the microscope through a liquid light guide (Thorlabs) and the power was determined to be 258.2 mW for 380 nm light and 203 mW for 470 nm light at the output of the light guide, respectively (15). The external patch clamp solution for Na_v_1.5 current recordings contained (in mM): 145 NaCl, 0.5 CdCl_2_, 2 CaCl_2_, 5 HEPES, 5 glucose and pH was adjusted to 7.4. The internal patch clamp solution for Na_v_1.5 current recordings contained (in mM): 30 NaCl, 100 CsCl, 10 HEPES, 2 MgCl_2_, 1 CaCl_2_, 2 MgATP, 0.05 NaGTP, 10 EGTA and 5 glucose the pH was adjusted to 7.4 using CsOH. AB2 was dissolved in external recording solution at indicated concentrations.

The patch clamp protocol to record Na_v_1.5 currents in the presence of AB2 depolarized the membrane potential from -70 mV to -10 mV for 100 ms at a frequency of 1 Hz. The recording cycle included 16 depolarizations under either 385 nm or 470 nm light illumination. The resulting Na_v_1.5 peak currents were analyzed and plotted. I-V relationships were elicited by stepping from -70 mV to potentials between -70 mV and +40 mV for 100 ms under simultaneous illumination with either 385 nm or 470 nm irradiation at a frequency of 1 Hz.

The I-V relationships were analyzed using Igor Pro 8 (WaveMetrics Inc., Portland, OR, USA). I-V curves were fit with I = G(V_m_-V_rev_)/(1 + exp[(V_m_-V_1/2)_/k)], where V_rev_ is the extrapolated reversal potential. A depolarized holding potential of -70 mV was chosen to exploit the high affinity of the bupivacaine derivative AB2 for inactivated Nav1.5 channels [6, 73] and simultaneously minimize interference of its use-dependency with photoswitching.

Quantification of Na_v_1.5 photoswitching (ps) was performed as described before [17]. Here, the peak inward Na_v_1.5 current of the 16^th^ depolarization at 380 nm or 470 nm illumination of two consecutive recorded cycles (one at 385 nm and one at 470 nm light) were analyzed: photoswitching = (I_380_ – I_480_/ I_380_) × 100; The resulting ps value represents only a qualitative indicator of light-induced block since AB2 was present all time. Data were analyzed in Igor Pro using the Patcheŕs Power Tools (MPI for Biophysical Chemistry, Göttingen) plugin.

### Two-electrode voltage clamp

Recordings were performed using a TURBO TEC-03X amplifier (npi electronic, Tamm, Germany). The glass capillaries (Science Products, Hofheim, Germany) were pulled to recording pipettes with tip resistances of 0.1 - 1 MΩ using a Micropipette puller (DMZ-Universal, Zeitz, Martinsried, Germany). The pipettes were backfilled with 3 mM KCl. Recordings of the oocytes were performed at room temperature in normal frog ringer solution containing (in mM): 115 NaCl, 2.5 KCl, 1.8 CaCl2 and 10 HEPES. The pH was adjusted to pH 7.2 by NaOH.

Use dependence block experiments and IC_50_ determination of lidocaine, bupivacaine, and *cis*-AB2 and *trans*-AB2 on Na_v_1.5 channel currents in *Xenopus laevis* oocytes were performed as described below. The holding potential was set to -100 mV. Pulse protocols were applied in a series of frequency intervals of 1, 5, 10, and 20 Hz at a duration of 10 s each. Before depolarization, the membrane was hyperpolarized to -120 mV for 20 ms or in case of the 20 Hz protocol for 10 ms respectively. Afterwards, the membrane was depolarized to +20 mV for 20 ms followed by a repolarization back to -100 mV. First, control recordings (w/o drug) were performed. After 3 min of incubation, the same experiments in the presence of the respective Na_v_1.5 blocker at a distinct concentration followed. For AB2, the duration at each frequency was doubled to 20 s, whereas half of the time the sample was illuminated at either 385 nm or 480 nm light yielding *cis*-AB2 and *trans*-AB2, respectively. Here, the above mentioned custom-built light source was applied and the light guide was positioned above the oocytes within the recording chamber. The resulting Na_v_1.5 peak currents were analyzed and plotted using Igor Pro 8 (WaveMetrics Inc., Portland, OR, USA) and GraphPad Prism 8 (GraphPad Software, Inc., La Jolla, CA, USA). The IC_50_ values were calculated using a nonlinear regression curve (equation 1) to fit steady-state currents of 1, 5, 10, and 20 Hz.

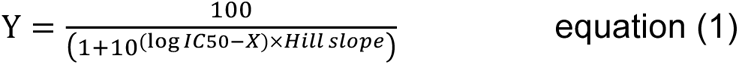

### Tissue slice preparation

Mouse hearts were perfused for 2 – 3 minutes on a Langendorff setup via coronary circulation with bicarbonate-buffered extracellular solution containing (in mM): 123 NaCl, 1.8 CaCl_2_, 5.4 KCl, 1.2 MgCl_2_, 1.4 NaH_2_PO_4_, 24 NaHCO_3_, and 10 glucose (pH 7.4) to ensure that isolated hearts beat rhythmically. Perfusion was subsequently changed to bicarbonate-buffered extracellular solution supplemented with the voltage-sensitive dye Di-8-ANEPPS at 3 µM (D3167, Invitrogen). Following dye loading for 4 minutes, hearts were perfused with 10 mM 2,3-butanedione monoxime (BDM) (B0753, Sigma) dissolved in bicarbonate-buffered solution until they ceased beating (∼ 5 min); they were then embedded in low-melt agarose (4 % in phosphate buffered saline) at 37 °C. The blocks were rapidly chilled until the agarose completely solidified. The agarose-embedded hearts were glued with tissue adhesive histoacryl (175182, Bbraun) on a specimen holder. Samples were sectioned transversally into 350 µm thick slices employing a vibratome (LEICA VT1200S) with steel blades at a progression speed of 0.03 mm/s and amplitude of 2 mm [68]. During the sectioning process, the tissue was kept in an ice-cold oxygenated bicarbonate-buffered solution (95 % O_2_, 5 % CO_2_) containing 10 mM BDM. The slice was immediately placed into a custom-made circulation chamber filled with 10 mM BDM- containing oxygenated bicarbonate-buffered solution (95 % O_2_, 5 % CO_2_). The temperature of the slice chamber was maintained at 35 °C by placing a glass spiral before the chamber entrance and controlling the solution flow rate through the chamber. A unipolar platinum-iridium electrode (ThermoFisher, Germany) which was coupled to an electrical stimulator (STG4002, Multi Channel Systems MCS GmbH) at a cycle duration of 200 ms, a pulse width of 1 ms, and 1.5 times the threshold strength was used to pace the slices (3 - 4 V). The cardiac slice was constantly paced for at least 30 min to recover, which was necessary to achieve an electrophysiological steady-state. During this time, the signal-to-noise ratio of fluorescence intensity also improved and achieved a steady-state [45, 68].

### Optical mapping

In order to excite Di-8-ANEPPS, a LED light source (LEX3-G; 525 nm; SciMedia/Brainvision) coupled with a THT Macroscope (SciMedia/Brainvision) was used. The light passed through an excitation filter (BP531 nm/40), was deflected by a dichroic mirror (580-FDI) towards the perfusion chamber, and was focused onto the slice. A 1.0X objective (Leica) was used to gather the fluorescence signals, which were then filtered using a long-pass filter (LP600 nm) and collected by a MiCAM05-N256 imaging system (SciMedia/Brainvision) equipped with a CMOS image sensor (mapping field area: 10 × 10 mm; frame rates: 1 kHz; SciMedia/Brainvision). The proprietary BV Workbench (Brainvision) was used to operate the optical recordings, and recordings were stored in the appropriate data format before being submitted to specialized MATLAB analysis.

After fluorescence reached a steady state, measurements were conducted over a 30-minute period in the absence of AB2 under dark conditions. Subsequently, 5 µM AB2 was directly introduced into the system, followed by a 30-minute direct exposure of the tissue slice to UV light at 385 nm at an approximate intensity of 16 mW/mm². This UV illumination induced the formation of *cis*-AB2, during which fluorescence data were collected. Finally, a transition from UV to blue light (470 nm) at the same intensity was employed to convert cis-AB2 to trans- AB2, with data acquisition at the end of this 30 minutes illumination period.

### Analysis of optical data

Data processing, analysis, and visualization were accomplished by custom-developed scripts written in MATLAB (ver. R2016b). Optical data processing included temporal stacking [64], drift correction by polynomial fitting, and 3 × 3 pixels spatial mean filtering [23, 37]. The data were analyzed by calculating activation time, AP duration at 80 % of repolarization (APD_80_), and AP amplitudes. Local activation periods of optically recorded APs were quantified in ms and were equated with the moment when the AP upstroke reached 80 % of its total amplitude. In order to assess AP durations, we adapted the “findpeaks” program to identify local maxima and widths at various peak amplitude levels. The amplitude of an AP was quantified as relative fluorescence intensity change 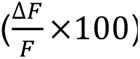. The vector drawing program Inkscape was used to rearrange and reduce MATLAB displays and maps [30].

### *Ex vivo* experiments in the intact heart

25-28 week old female CD1 wild type mice were sacrificed by cervical dislocation. Afterwards, hearts were explanted and treated with a heparin containing Tyrode’s solution at 4 °C. In the following step, hearts were cannulated in the Langendorff configuration (17). Here, the coronary arteries were retrogradely perfused with Tyrode’s solution at 33-35 °C. The solution contained (in mM): 140 NaCl, 5.4 KCl, 1.8 CaCl_2_, 2 MgCl_2_, 10 D-glucose and 10 HEPES (pH 7.4, adjusted with NaOH). *Cis*-AB2 and *trans*-AB2 were applied within the perfusion solution to the hearts. *Cis*- and *trans*-AB2 states were set by pre-irradiation with either 380 nm light or 480 nm light in a custom-made perfusion chamber combined with a custom-built light source as described above [17]. LED power at the entrance of the perfusion chamber was determined to be 258.2 mW for 380 nm light and 203 mW for 470 nm light, respectively.

For the detection of native field potentials (FPs), bipolar ventricular cardiac electrograms were recorded by two Ag-electrodes from the left ventricle. The latency of the FP was detected as the interval between the stimulus and the peak repolarization (Fig. 3f). Furthermore, we determined the murine native field potential width (FPW) by measuring the peak width at 90% of its height. To assess the drug effect in its time course, we calculated the linear slope of FPW changes over time for control, cis-AB2, and trans-AB2 conditions.

Electrical stimulation of the heart to determine ventricular field potentials were either performed as constant pacing experiments at 400 bpm (S1-S1 = 150 ms, 6.6 Hz) or as S1- S2 protocols. Here, stimulation electrodes were positioned at the right ventricle and perpendicularly located bipolar detection electrodes at the left ventricle. For constant pacing experiments the FP latency and width were determined as defined in Fig. 2f. S1-S2 protocols included looped intervals. Six S1 intervals with a duration of S1-S1 = 150 ms (400 bpm) were followed by an S1-S2 interval. Here, interval length is shortened every loop by either 1, 3, or 5 ms starting from 150 ms. At the end of the loop, an S2 interval of 150 ms followed. The resulting FPs within the S2 interval were analyzed for their latency. The first electrical stimulus that failed to produce a FP represented the effective refractory period (ERP), which was correlated to a distinct frequency. We designed drug application sequences as protocols A – C. Sequence A comprised 5 minutes control (w/o AB2) and 7 minutes of *cis*-AB2 (Fig. 4b). Protocol B included 5 minutes control (w/o AB2), 1 minute of *cis*-AB2, and 6 minutes of *trans*-AB2. Protocol C consisted of 5 minutes control, 4 minutes of *cis*-AB2, and 3 min of *trans*-AB2. Changes in the conduction velocity (CV) were determined in S1 intervals by comparing FP latencies under control, *cis*-AB2 and *trans*-AB2 conditions. Here, a decrease in CV is defined as (latency_cis/trans_ / latency_control_) × 100. All data related to protocol A – C were recorded in low K^+^ Tyrode’s solution (in mM: 140 NaCl, 2 KCl, 1.8 CaCl_2_, 2 MgCl_2_, 10 D-glucose and 10 HEPES (pH 7.4, adjusted with NaOH). Ventricular arrhythmias (VA), were either triggered in the presence of a pro-arrhythmic low K^+^ Tyrode’s solution (see above) by applying electrical burst stimulation, S1-S2 protocols, or occurred spontaneously. Burst stimulation included 80 stimuli at 50 Hz for 1.6 s with 5 V pulse amplitude. A VA was considered to be stable after a continuous duration of at least 5 minutes. VAs were detected in a time window of 12 minutes and drugs were applied according to protocols A-C.

For the detection of signals, a bio-amplifier recording system was used (PowerLab 8/30, Animal Bio Amp ML 136, LabChart 7.1 software, AD Instruments). Analysis was performed with custom-built Matlab codes.

### Heat map

To illustrate the recorded data of VA in a comprehensive way, we chose the format of a heat map. Individual experiments resulting from control condition (w/o AB2) and protocols A-B were divided into intervals of 10 s. For each interval, a Fourier analysis was performed resulting in a frequency spectrum, which was normalized to a total integral of 1.

Consecutive time intervals of normalized and averaged VA frequencies spectra under control conditions or protocol A or B formed the temporal course of the heat map. For quantification of the frequency changes, we determined the weighted frequency of the first and the last minute of a heat map and indicated the results in the same representation (Fig. 5 j-l).

### Synthesis

Chemical synthesis of AB2 was carried out as described before [39]

### Mathematical Modelling

The Bondarenko model was used as a single cell model of mouse left ventricle AP [8]. We replaced the Markov model of sodium channels by a corresponding Hodgkin-Huxley model to prevent observed repolarization failure in 2D simulations [43]. The model was paced at 150 ms cycle length to reach a steady state. Inhibitory effects of AB2 on mouse APs were modeled by simulation of all combinations of g_Na_, g_Kr_, g_Kur_, g_Kto_ reductions (80%, 60%, 40% and 20%). For VA simulation on 2D tissue, a 1.2 cm rectangle of ventricular tissue with periodic boundaries was modeled and the extracellular K^+^ concentration of 2 mM was included to resemble Langendorff experimental setups. A sustained VA was induced by a re-entry initiation with a cross-shock protocol. The first stimulus applied at the right of the sheet was followed by a premature stimulus applied on a square area at a subsequent time [13]. I_Na_ and I_Kur_ block were modeled by 60 % reduction of corresponding conductances (g_Na_, g_Kur_). The core area of generator spirals was calculated by AP analysis of each node identifying the time of the peak with minimum prominence and < 15 mV. AP analysis was performed using MATLAB. All simulations were implemented in the openCARP environment [52].

### Statistical analysis

All data in the present publication are expressed as mean ± standard error of the mean (s.e.m.). Unless otherwise stated, data were analyzed using paired Student’s t test or cox regression analysis. Asterisks indicate levels of statistical significance with * p ≤ 0.05, ** p ≤ 0.01, and *** p ≤ 0.001.

### Data availability

Data generated and analyzed in this study are included in this article or may be obtained by the corresponding authors on reasonable request.

## Supporting Information (SI)

### IC_50_ validation

To validate the determined IC_50_ values of AB2 at different concentrations we additionally determined the IC_50_ of bupivacaine on hNa_v_1.5, resulting in resting state IC_50_ of 81.56 ± 8.3 µM at 1 Hz, of 34.16 µM ± 4.9 µM at 5 Hz, of 19.63 ± 3.4 µM at 10 Hz, and of 7.15 ± 1.5 µM at 20 Hz, respectively (n = 57 oocytes). For 1 Hz, these values are in close proximity to the ones reported in the literature: [2] IC_50_ = 69.5 µM (HEK293, outside-out), [1]: IC_50_ = 50 µM (HEK293, whole cell), underlining the reliability of our recorded data. The apparent affinity of AB2 for hNa_v_1.5 is hence only slightly lower compared to its parent molecule bupivacaine.

**Fig. S1.**
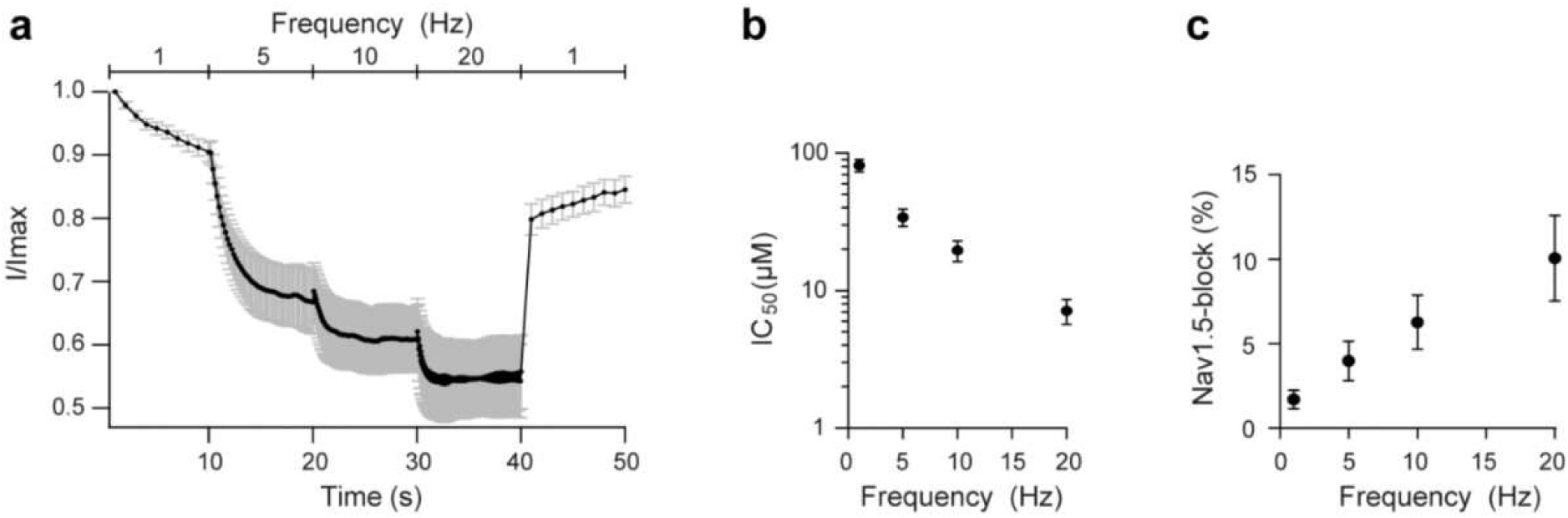
IC_50_ validation for bupivacaine on hNav1.5 in *Xenopus* oocytes. **a** Normalized hNav1.5 peak currents demonstrate the use-dependency of bupivacaine (50 µM, n = 10 oocytes), hNav1.5 was activated at frequencies of 1, 5, 10, and 20 Hz, respectively. **b** Use-dependency of the IC_50_ of bupivacaine (logarithmic representation of IC_50_, n = 57 oocytes). **c** Relative block of hNa_v_1.5 current by 1 µM bupivacaine increases with activation rate. All data expressed as mean ± s.e.m.

**Fig. S2.**
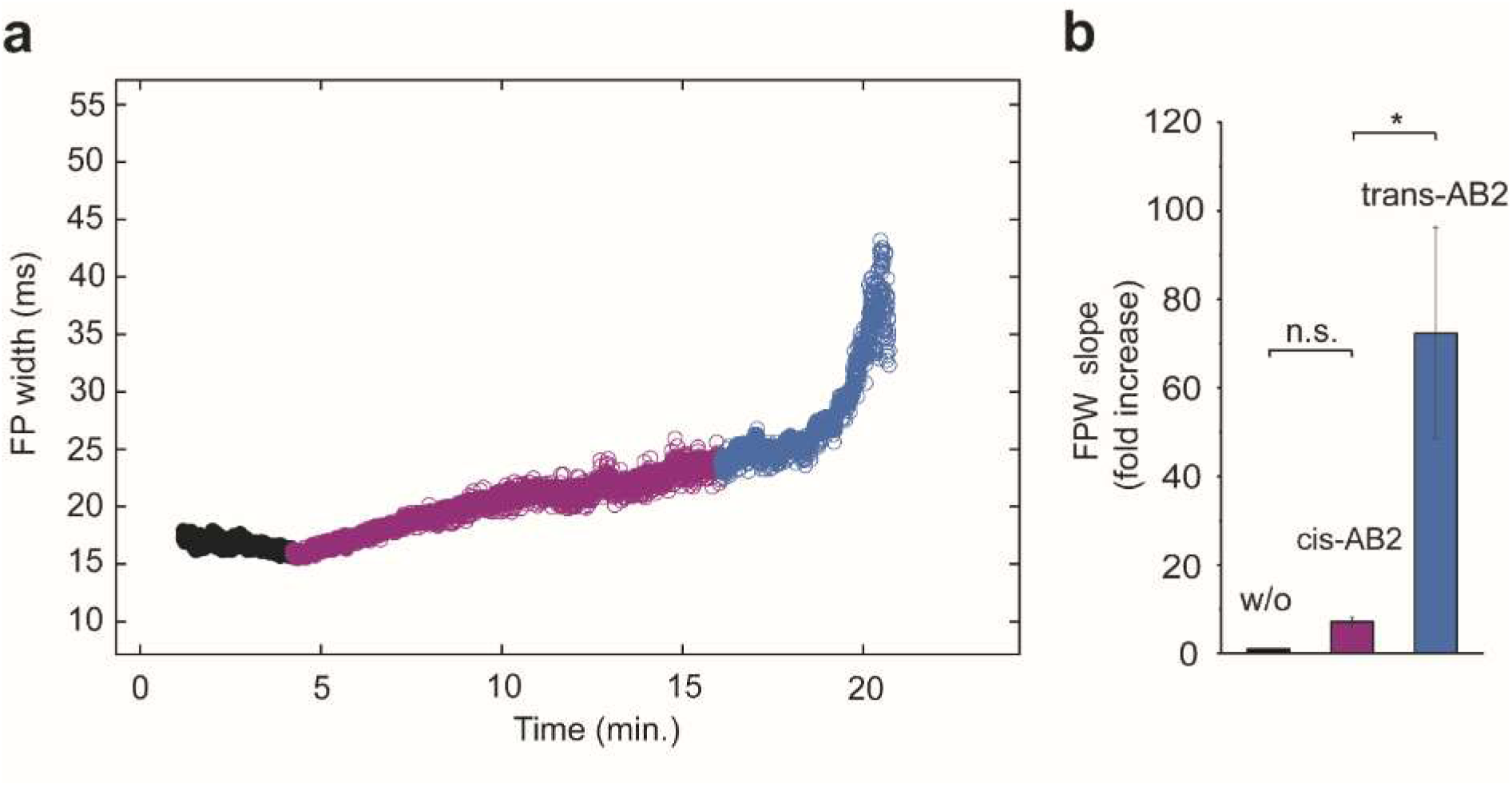
Stimulated field potential width under control, 10 µM *cis*-AB2 and 10 µM *trans*-AB2 in a Langendorff-perfused mouse heart preparation. a Time course of field potential width development under indicated conditions and **b** its quantification (paired t-test, w/o AB2 to *cis-*AB2 p=0.0524, *cis*-AB2 to *trans*-AB2 p=0.0044, n = 3 hearts). n.s. = not significant, * p < 0.05, All data expressed as mean ± s.e.m .

**Fig. S3.**
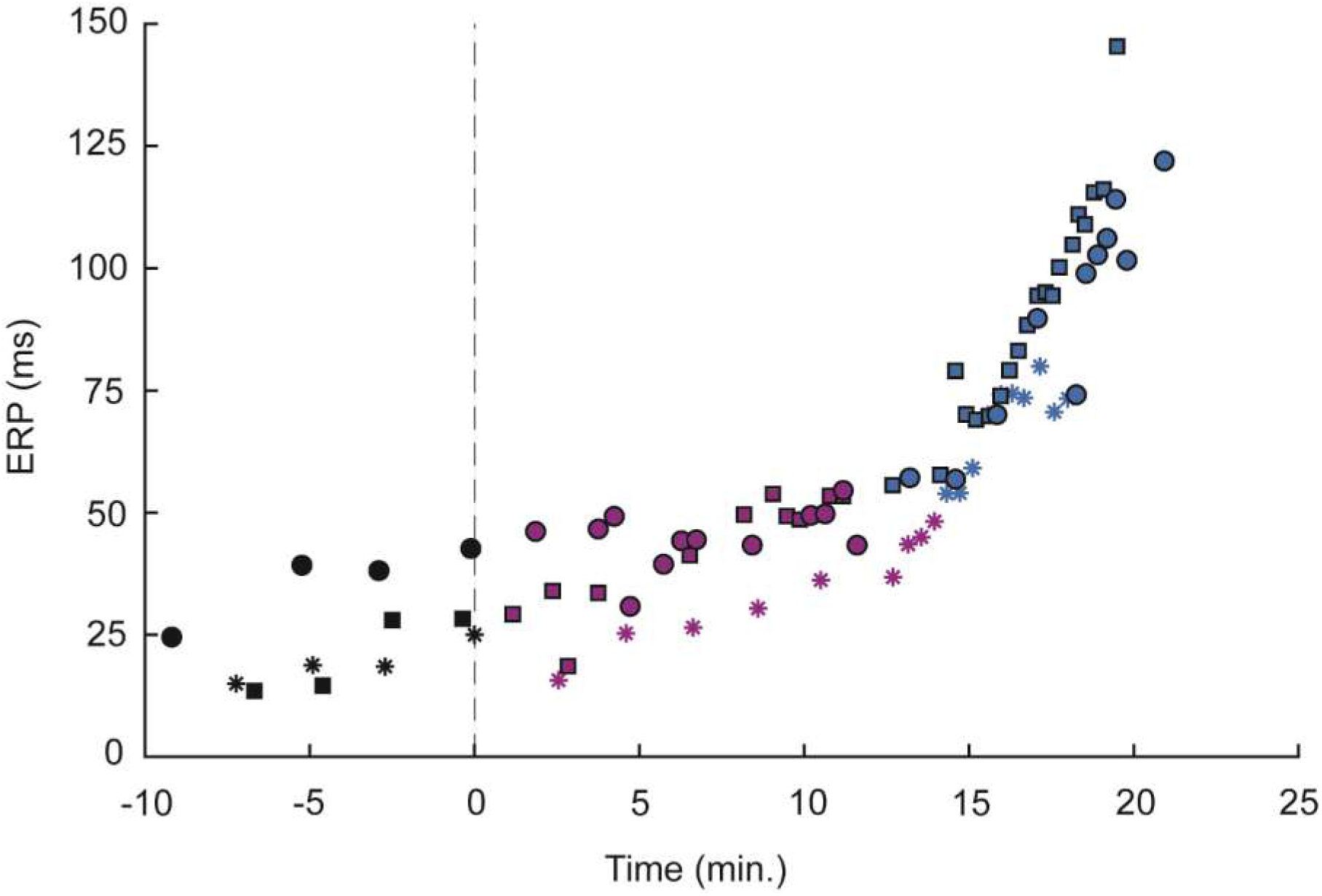
ERP time course under control, 5 µM *cis*-AB2 and 5 µM *trans*-AB2 conditions (n = 3 hearts).

**Fig. S4.**
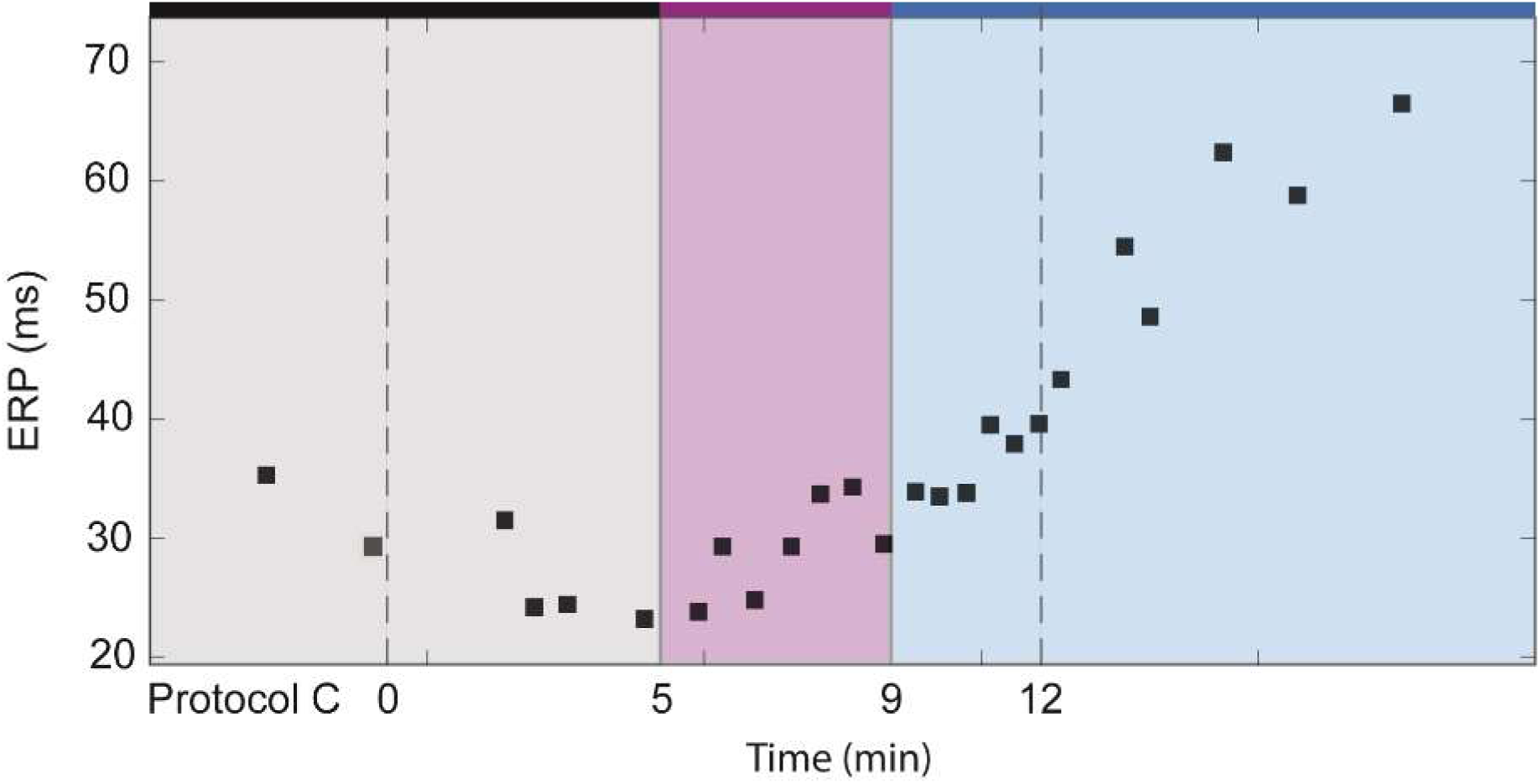
Representative ERP time course under control, 5 µM *cis*-AB2 and 5 µM *trans*-AB2 conditions according to protocol C (n = 3 hearts).

**Fig. S5.**
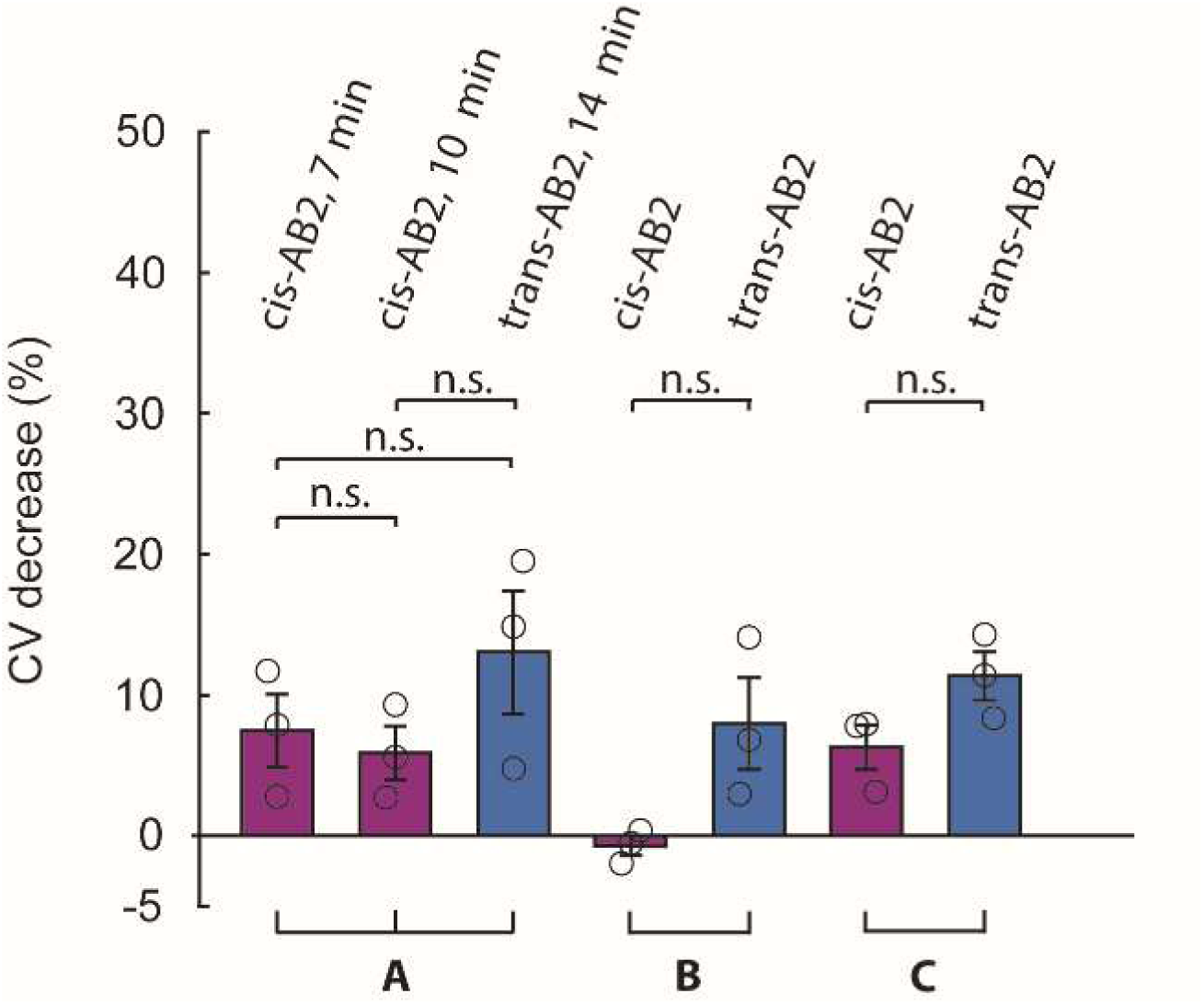
Relative CV decrease at a basic cycle length of 150 ms in field potential experiments. Quantification of normalized CV decrease under *cis*- and *trans*-AB2 according to protocols A to C. Statistical analysis: paired sample Students’ t-test (n = 3), *cis-*AB2 (7 min) to *cis-*AB2 (10 min) (protocol A) p = 0.1725, *cis-*AB2 (7 min) to *trans-*AB2 (14 min) (protocol A) p = 0.4403, *cis-*AB2 (10 min) to *trans-* AB2 (14 min) p = 0.2984, *cis-*AB2 to *trans-*AB2 (protocol B) p = 0.13, *cis-*AB2 to *trans-*AB2 (protocol C) p = 0.2507, n.s. = not significant. All data expressed as mean ± s.e.m.

**Fig. S6.**
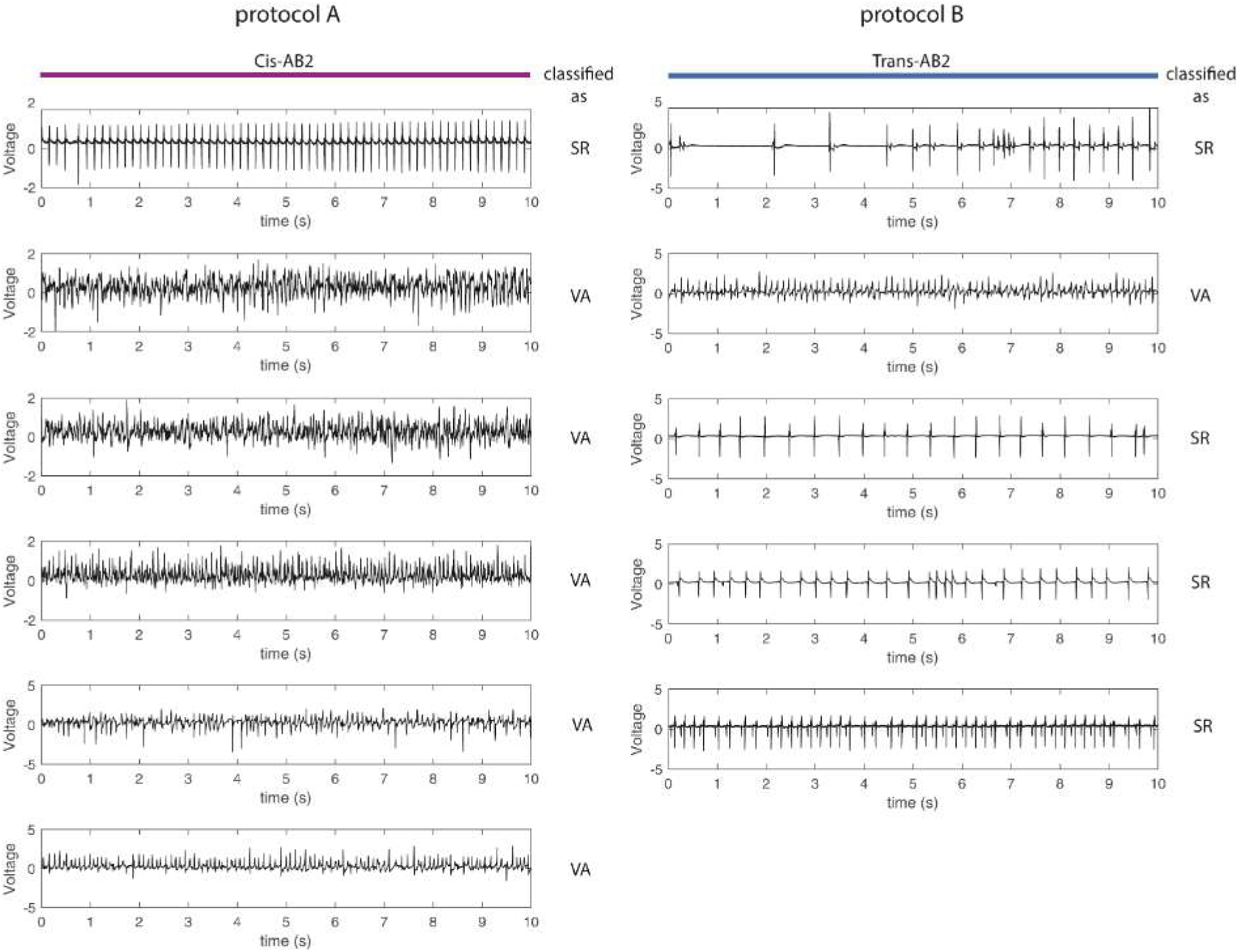
Termination rate of VA. Individual field potential recordings between 00:11:40 - 00:11:50 under *cis*-AB2 (protocol A, n = 6 hearts) and *trans-AB2* (protocol B, n = 5 hearts).

**Fig. S7.**
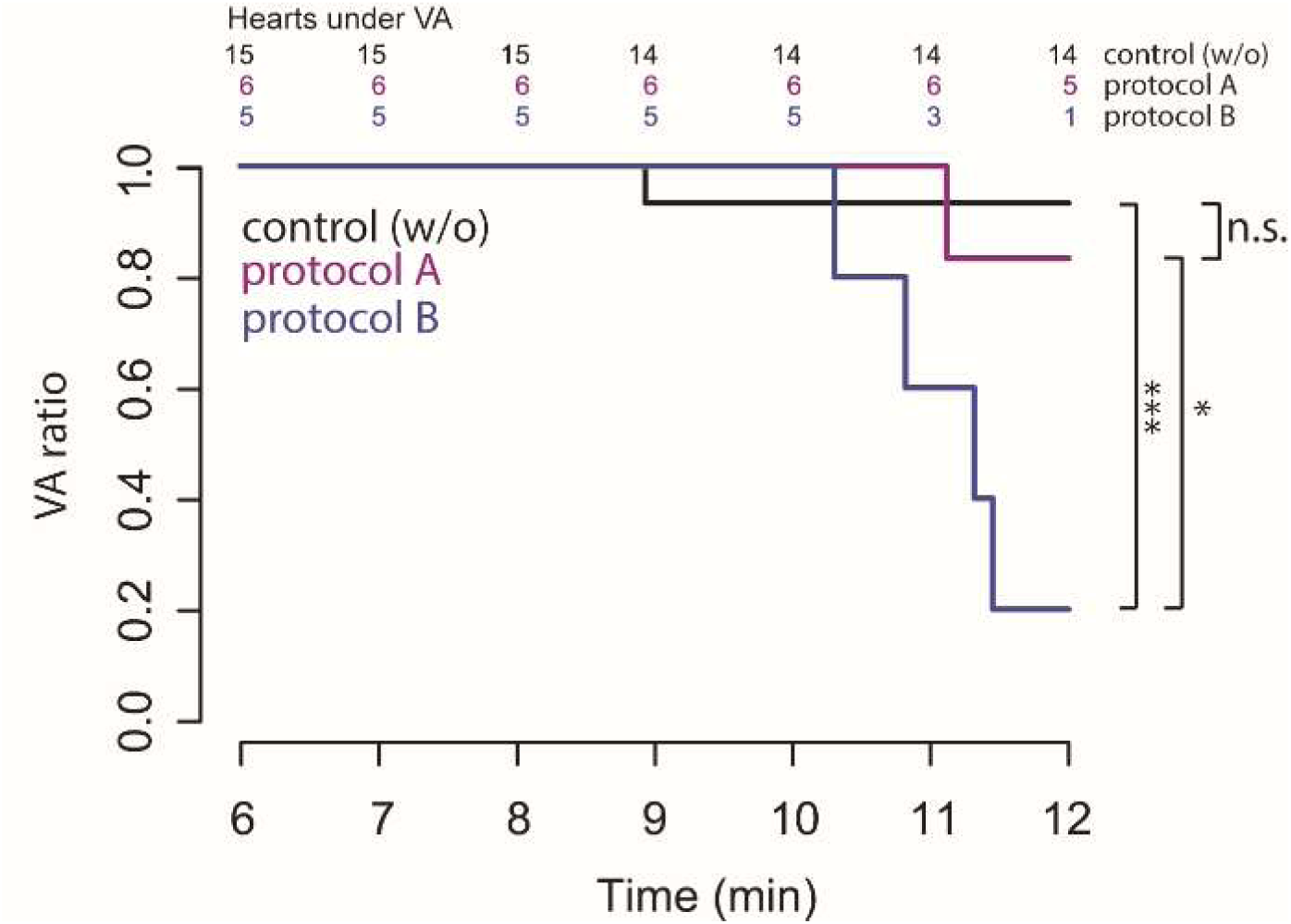
Kaplan Meier plot of VA termination by *trans*-AB2. VA ratio is plotted over time, representing the time interval of protocols A-B and control situation (6-12 min). Statistical analysis for the period 6- 12 min: control to *cis*-AB2 (protocol A) Hazard ratio = 0.41, p = 0.5, *cis*-AB2 (protocol A) to *trans*-AB2 (protocol B) Hazard ratio = 4.0, p = 0.04, control to *trans*-AB2 (protocol B) Hazard ratio = 10.8, p = 0.001. Univariate Cox proportional hazards regression model was used (coxph package, R), n.s. = not significant, * p ≤ 0.05, *** p ≤ 0.001. All data expressed as mean ± s.e.m.

**Fig. S8.**
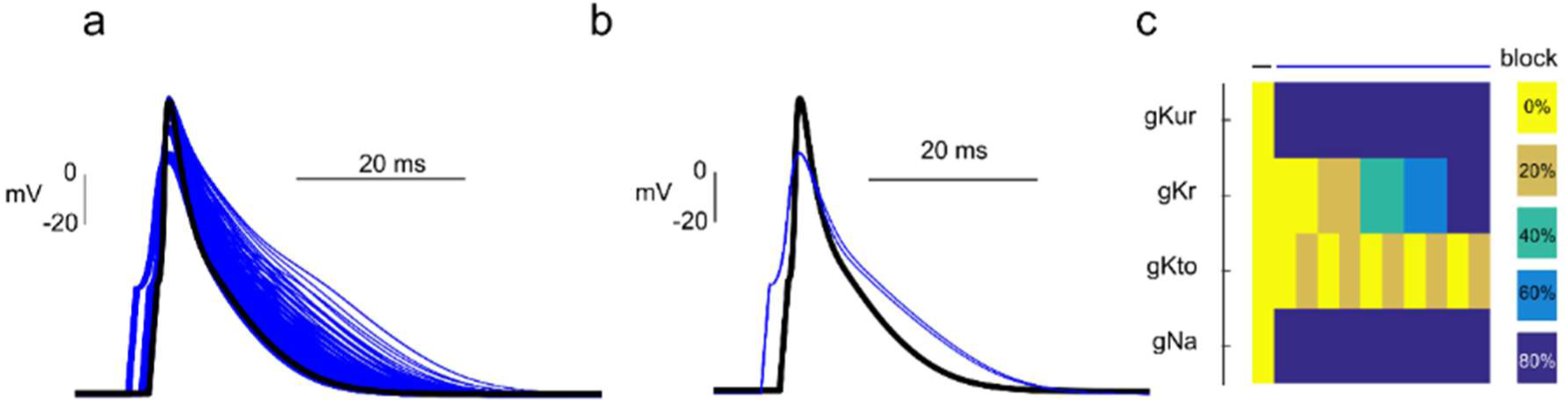
Simulation of single cardiomyocyte action potentials under combinatorial inhibition of Na^+^ and K^+^ conductivities. **a** Overlay of a normal action potential (AP) and APs after combinatorial inhibition of g_Kur_, g_Kr_, g_Kto_ and g_Nav_ (black and blue, respectively) by increasing steps of 20 %. **b** APs filtered for shortest APD_10_, longest APD_80_, and maximum reduction in peak amplitude (blue) overlayed on control (black). **c** Percentage of inhibition of indicated ionic conductivities yielding the filtered APs in (b).

**Fig. S9.**
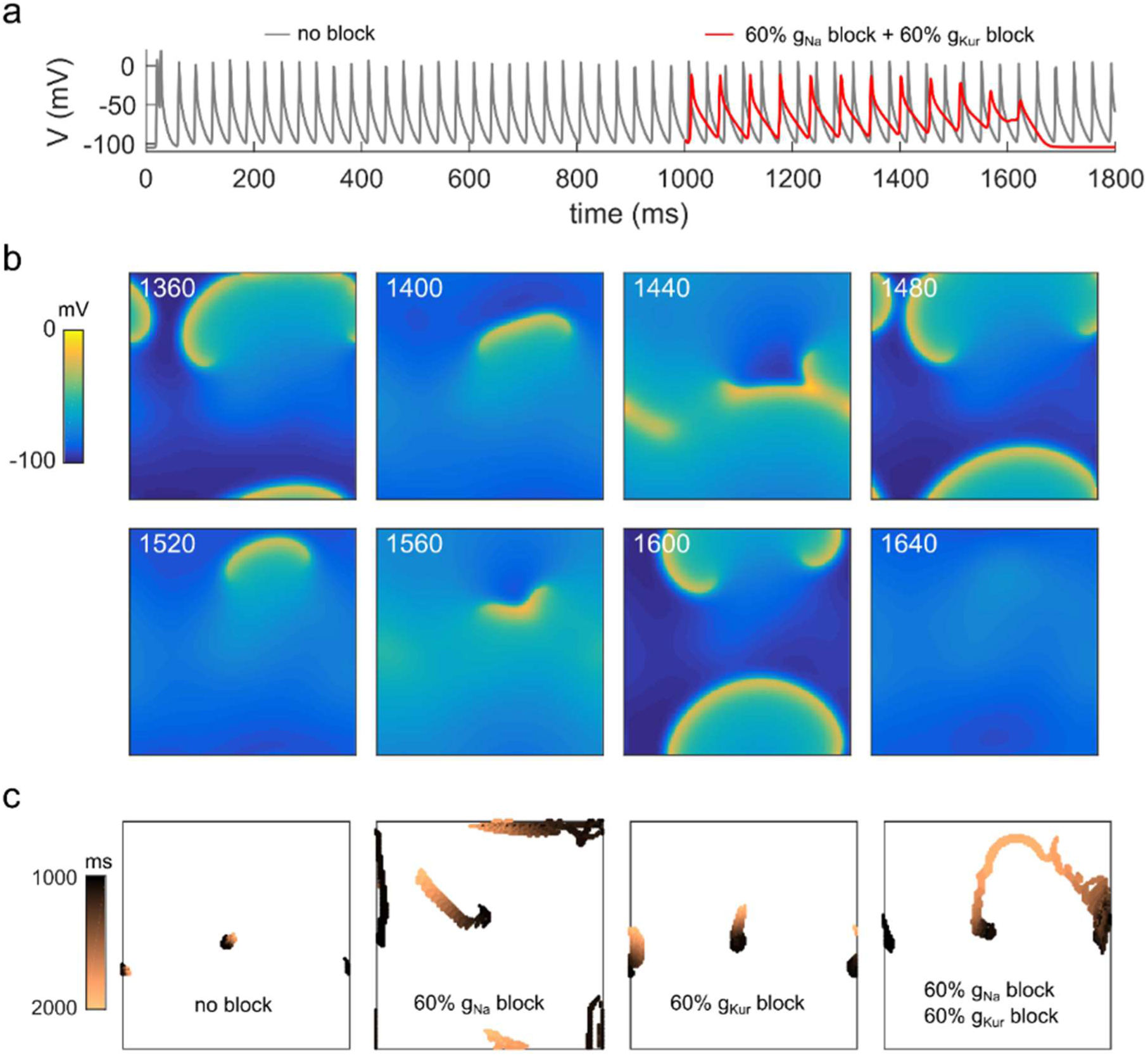
Termination of sustained arrhythmia in a two-dimensional model of cardiac tissue by combined Na^+^ and K^+^ conductivity block. **a** Sustained electrical fibrillation is terminated by a combined reduction of g_Nav_ and g_Kur_ by 60 % each. **b** Snapshots of spatially resolved transmembrane voltage at indicated simulation time points after combined reduction of g_Nav_ and g_Kur_ by 60 % each. **c** Trajectories of spiral core tips of simulated sustained arrhythmia were calculated for the indicated experimental conditions. Note the increased meandering of core tips eventually leading to the collision of trajectories and subsequent termination of re-entries after combined reduction of g_Nav_ and g_Kur_ by 60 % each.

## Notes

### Competing Interest Statement

The authors have declared no competing interest.

